# Evaluating Inositol phospholipid interactions with Inward Rectifier Potassium Channels and characterising their role in Disease

**DOI:** 10.1101/2020.09.03.281378

**Authors:** Tanadet Pipatpolkai, Robin A. Corey, Peter Proks, Frances M. Ashcroft, Phillip J. Stansfeld

## Abstract

Membrane proteins are frequently modulated by specific protein-lipid interactions. The activation of human inward rectifying potassium (hKir) channels by phosphoinositides (PI) has been well characterised. Here, we apply a coarse-grained molecular dynamics free-energy perturbation (CG-FEP) protocol to capture the energetics of binding of PI lipids to hKir channels. By using either a single- or multi-step approach, we establish a consistent value for the binding of PIP_2_ to hKir channels, relative to the binding of the bulk phosphatidylcholine phospholipid. Furthermore, by perturbing amino acid side chains on hKir6.2, we show that the neonatal diabetes mutation E179K increases PIP_2_ affinity, while the congenital hyperinsulinism mutation K67N results in a reduced affinity. We show good agreement with electrophysiological data where E179K exhibits a reduction in neomycin sensitivity, implying that PIP_2_ binds more tightly E179K channels. This illustrates the application of CG-FEP to compare affinities between lipid species, and for annotating amino acid residues.

## Introduction

Ion channels are integral membrane proteins that mediate ionic flux across the plasma membrane. This process can be regulated by the binding of factors such as soluble ligands or phospholipids to the channel. In particular, lipid binding has been shown to affect many types of ion channel, regulating both their oligomeric state and their activation^1^. Impairment of these processes can lead to a range of human and animal diseases. One well-studied class of ion channels are the mammalian inward rectifying potassium (Kir) channels. In the case of Kir6.2, the pore component of the ATP-sensitive potassium (K_ATP_) channel complex, mutations may result in either a loss, or gain, of channel function, resulting in congenital hyperinsulinism (CHI) and neonatal diabetes (NDM), respectively^7^.

Kir channels are activated by phosphoinositides, in particular phosphatidylinositol-4,5-bisphosphate (PIP_2_)^8^. Different Kir channels exhibit variable affinities and levels of channel activation to different phosphoinositides^9–11^. The PIP_2_ binding site on Kir channels has been well defined in several crystal structures, such as chicken Kir2.2 [PDB entry: 3SPI]^12^ and mouse Kir3.2 [PDB entry: 3SYA]^13^. Meanwhile, recent advances in cryo-electron microscopy (cryo-EM) have enabled several high-resolution structures of the pancreatic K_ATP_ channel complex (Supplementary Table 1), which comprises a central tetrameric pore formed of Kir6.2 subunits, surrounded by four regulatory sulfonylurea receptor 1 (SUR1) subunits^14^. This octameric complex couples pancreatic beta-cell energy status to insulin secretion^15^. Mutations in the Kir6.2 subunit that are located near the PIP_2_ binding site are associated with NDM (e.g. E179K/A) and CHI (e.g. K67N)^16–18^. A previous study has shown that the K67N mutation does not alter channel surface expression but has reduced channel activation when cell metabolism was inhibited^18^. The mechanism of how E179K/A and K67N mutations affect channel activity is currently unclear.

Kir channels form an attractive target for applying a computational approach to compare binding affinity between phosphoinositides and also assess the impact of mutations on binding. Coarse-grained (CG) molecular dynamics (MD) simulations have previously been used to identify lipid-binding sites on ion channels^3–5^, as well as predicting the affinity of interactions^6^.

For many years, the application of atomistic free energy perturbation (FEP) methods have been successfully applied to determine small molecule, lipid, and drug binding affinities^19^ as well as to study the impact of amino acid side chain mutations^20,21^. Our recent study showed how the method could be extended to a CG protocol (CG-FEP) to assess relative protein-lipid binding free energies. This approach was in strong agreement with other free energy calculation methods such as potential of mean force calculation (PMF) and well-tempered metadynamics (WTMetaD)^6^.

In this study, we use CG-FEP^6^ to compare the relative binding free energies between different phospholipids and the human Kir6.2 channel, capturing the full thermodynamic cycle for the transition of PIP_2_ to PC, either directly or via intermediates phosphatidylinositol-4-phosphate (PI4P) and phosphatidylinositol (PI), and thereby reporting on the affinity of each interaction. We extend the methodology to investigate the functional effect of lipid-associated neonatal diabetes mutations (E179K/A) and a congenital hyperinsulinism (CHI) mutation (K67N) in hKir6.2^22,23^. Based on the predicted binding site for PIP_2_, we calculate that these residues interact with PIP_2_ in the membrane. This therefore provides a biochemical and structural explanation for the different clinical phenotypes.

We couple these analyses with electrophysiology, to assess both the affinity for, and channel activation by, PIP_2_. We also extend the computational methodology to assess the binding free energy differences between a range of hKir channels (hKir1.1, hKir2.2, and hKir3.2) and other inositide lipid species such as phosphatidylinositol-4-phosphate (PI4P) and phosphatidylinositol (PI). Together, our application of the CG-FEP describes the affinity of membrane proteins with a range of different lipids, as well as examining how biologically important mutations affect these interactions.

## Results

### PIP_2_ binding conformation to the hKir6.2 channel

The initial position of the PIP_2_ molecule was obtained from the chicken Kir2.2 channel:diC8-PIP_2_ complex^12^. After structural alignment of hKir6.2 with chicken Kir2.2, one of the bound diC8-PIP_2_ molecules was extracted, converted to CG and the resultant hKir6.2-PIP_2_ complex was built into a PC membrane and simulated for 1 μs (n = 5) using CG lipid self-assembly^3,4,24^. We defined residues that were within a 6 Å radius of the whole PIP_2_ molecule for >75 % of simulation time as proximal residues (Fig. 1a). We found that PIP_2_ binds in the vicinity of both the N- and C-termini of the hKir6.2 channel, including ^67^KWP^69^, on the N-terminus, and the residues between 170 and 179 on the C-terminus. These regions contain a number of basic residues, which allows them to interact with the negatively charged phosphate groups on the inositol ring of the PIP_2_ headgroup. E179 is the only negatively charged amino acid to be within this cut-off from the lipid.

**Fig. 1.**
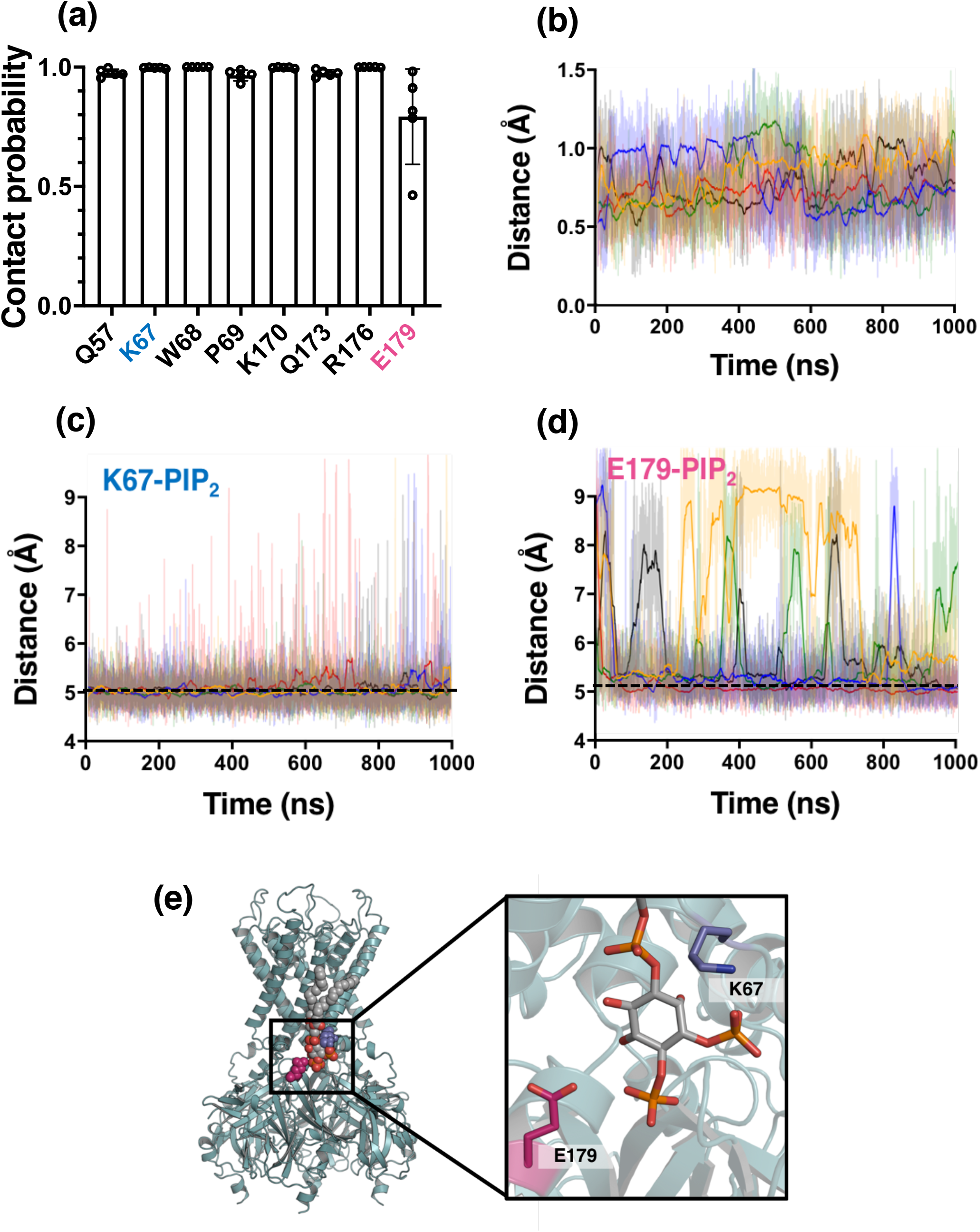
A PIP_2_ binding site on hKir6.2. **a** PIP_2_ contact analysis showing the fraction of time that residues are in 6 Å proximity to the PIP_2_ molecule (contact probability). Only residues with a >75% contact probability are shown. Data from 5 repeats of 1 μs simulations. **b** Root mean square deviation (RMSD) analysis of the PIP_2_ molecule headgroup when bound to the hKir6.2 tetramer. The different colours indicate the individual repeats of the simulations (n=5). The darker lines show the running average for each simulation. **c** Calculated distance between K67 and the PIP_2_ headgroup during a 1 μs simulation. The different colours indicate the individual repeats of the simulation. The darker line shows a running average. A dashed black line denotes the distance cutoff used to denote a contact in panel (a). **d** As in panel (c) but for E179 and the PIP_2_ headgroup. **e** PIP_2_ binding site on the hKir6.2 tetramer (green) showing PIP_2_ (grey with CPK colours), E179 (magenta) and K67 (blue).

We assessed the stability of PIP_2_ in its binding site using Root Mean Square Deviation (RMSD) analysis over the 1 μs simulations (Fig. 1b). The data show that the position of the PIP_2_ diverges very little in 1 μs (RMSD = *ca.* 0.8 Å). This was corroborated by analysing the distance between the PIP_2_ molecule and two amino acids near the PIP_2_ binding site, K67 and E179 (Figs. 1c and 1d). We found that the minimum distance between K67 and E179 and the PIP_2_ head group are approximately 5 Å respectively (Supplementary Fig. 1). Therefore, we hypothesise that mutations to these residues may affect the binding free energy of PIP_2_ to the channel.

### Step-wise perturbation of the PIP molecule bound to the hKir6.2 channel

We next assessed the contribution that each of the different PIP headgroup moieties (i.e. each phosphate group and the inositol ring) make to the free energy of binding to the closed hKir6.2 tetramer using CG-FEP. For this, we iteratively perturbed single beads to transform from one phospholipid (such as PIP_2_) into another (such as PC). This enabled us to calculate the binding free energy difference (ΔΔG) of the two different phospholipids to hKir6.2, embedded in a PC bilayer (Fig. 2a). For simplicity, all of our lipids have both palmitoyl and oleoyl alkyl chains. The energies were computed using Multistate Bennett Acceptance Ratio (MBAR)^25^, with convergence seen within *ca.* 200 ns per window (Supplementary Figs. 2a-b, 3a-b, 4-b and 5a-b). To prevent the molecule from leaving its binding site, we applied a flat-bottom distance restraint between the protein and lipid using Plumed^26,27^ (Supplementary Figs. 2a-c and 3a-c). This was mostly applicable for the calculations in which the lipid was transformed to PC. In the cases where the lipids remain bound at the binding site, applying a flat-bottom restraint makes no difference to the binding free energy and its convergence. (Supplementary Figs. 4c and 5c). This procedure also reduced the errors between simulation replicas. The infrequency with which the lipid experiences the restraint suggests that it has negligible effect on the binding energies (Supplementary Figs. 2d and 3d).

**Fig. 2.**
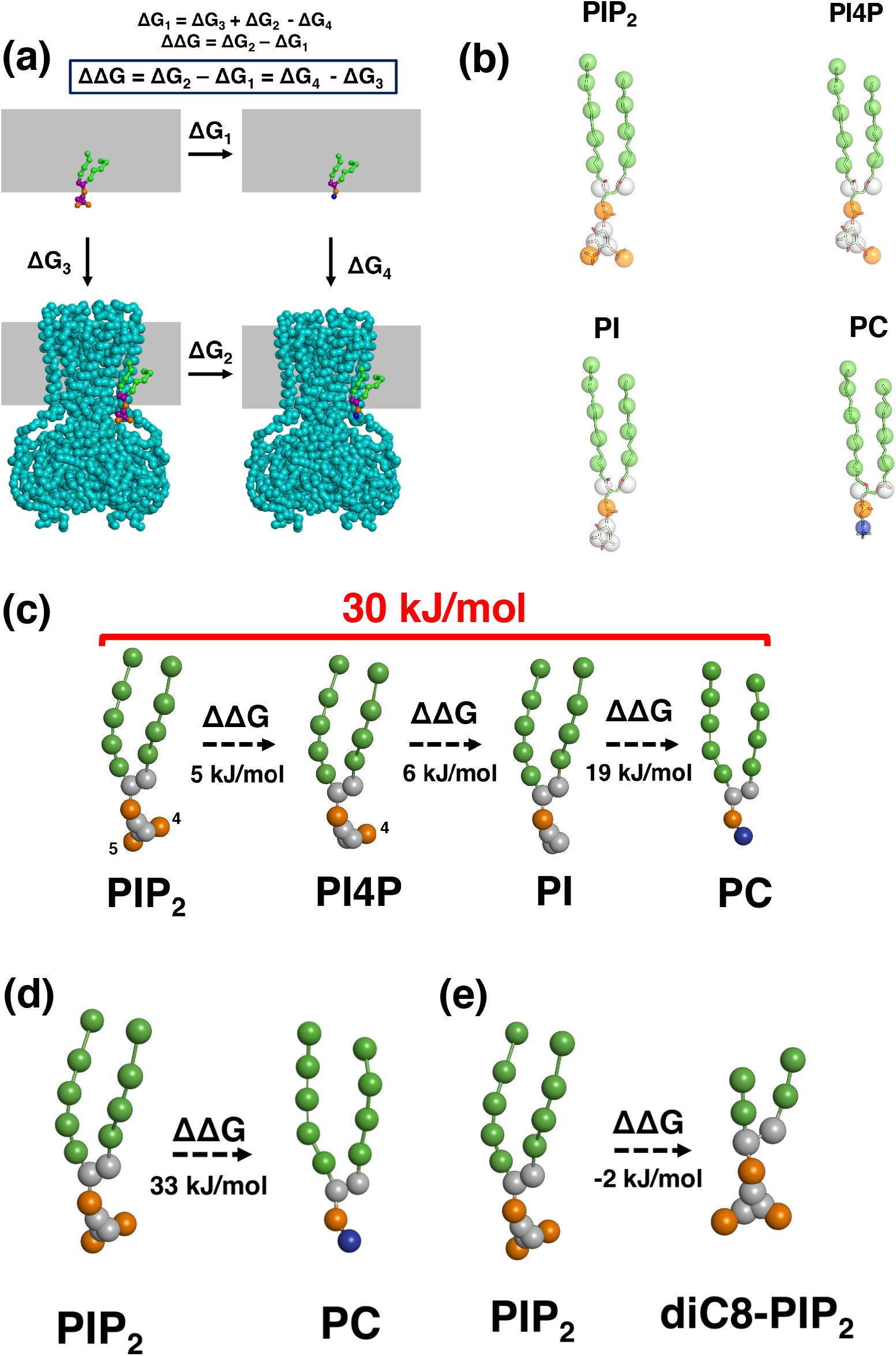
The free energy calculation of an individual phosphate group and fatty acid chains on a hKir6.2 tetramer. **a** Thermodynamic cycle used for the relative binding free energy calculations. The perturbation of the PIP_2_ headgroup (purple) was calculated in both the channel bound state (ΔG_2_) and free in the PC membrane (grey rectangle (ΔG_1_)). ΔG_3_ and ΔG_4_ can be calculated using such methods as PMF calculations (Figure 2 - Figure Supplement 6) **b** Coarse-grain to atomistic mapping of the phosphoniositides (PIP_2_, PI4P, PI and PC) **c** Change in binding free energy (ΔΔG) when individual phosphate groups are perturbed (i.e. from PIP_2_ to PI4P, from PI4P to PI and from PI to PC (values in black). The sum of these free energy changes (i.e. from PIP_2_ to PC) is given in red. Values are rounded to the nearest whole number. **d** Change in binding free energy when PIP_2_ is perturbed to PC. Values are rounded to the nearest whole number. **e** Change in binding free energy when PIP_2_ is perturbed to PIP_2_-diC8. Values are rounded to the nearest whole number

Transformation from PIP_2_ to PI4P showed a very small relative free energy change (Fig. 2b and Supplementary Table 2). This suggested that a phosphate group at either the 5’ position does not make a considerable contribution to PIP_2_ binding (Fig. 2b and Supplementary Table 2). However, we observed large free energy changes when the last phosphate group at the 4’ position and the inositol ring were perturbed (Fig. 2b and Supplementary Table 2). Summation of each individual moiety from PIP_2_ to PC gives a binding free energy of 30 ± 1 kJ/mol, which is remarkably similar to the 33 ± 3 kJ/mol we obtain for direct perturbation of PIP_2_ to PC (Fig. 2c, Supplementary Table 2, and Supplementary Fig. 6). This suggests that the approach we use is valid for both single- and multi-step free energy calculations.

A previous study demonstrated that crosstalk between different anionic lipids can affect the affinities of each lipid for the Kir2.2 channel^28^. Here, we show that the presence of 10% anionic lipid (POPS) in the lower leaflet of the bilayer does not affect the overall ΔΔG for PIP_2_ binding to Kir6.2 (Supplementary Figure 7).

Previous electrophysiological and crystallographic studies have commonly used the soluble eight-carbon atom phosphatidyl inositol, diC8-PIP_2_, to study channel activation^12,13,17^. Therefore, we investigated the effect of the length of the acyl chain on PIP_2_ binding affinity. We found that truncation of the acyl chain from either 4 or 5 particles (i.e. palmitoyl and oleoyl) to 2 particles (equivalent to 8 carbon atoms) had no effect on PIP_2_ affinity (Fig. 2d). This suggests that PIP_2_-diC8 is indeed an effective substitute for investigating the impact of PIP_2_ binding in electrophysiological and structural studies.

### Relative binding free energy calculations for PIP_2_ interactions with both wild-type and mutant hKir6.2

Based on the closed state model of Kir6.2 [PDB: 6BAA], we generated three structural models of hKir6.2 with disease-associated mutations; K67N which causes CHI, and E179A and E179K which cause neonatal diabetes^22,23^. As these residues are in close proximity to the PIP_2_ binding site (Fig. 1e), we hypothesised that mutations to these residues would modulate PIP_2_ affinity and thereby affect basal channel activity (i.e., the channel open probability, Po). An increase in PIP_2_ binding affinity should correlate with an increase in channel activation (Po) and thus also a reduced inhibition by ATP.

We next performed calculations in which the mutated hKir6.2 residue was perturbed in the presence and absence of PIP_2_ (Fig. 3a). This allows us to calculate the relative changes in the PIP_2_ binding free energy between the wild-type and mutant channels. For the highest energy residue substitution (E179K), we observed convergence of the free energy calculations within 50 ns per window (Supplementary Fig. 8). The data show an increase in PIP_2_ binding energy, and hence an increased affinity, with the E179K transformation. We also observed an increase in binding free energy with the E179A mutation (Fig. 3b and Supplementary Table 4). Conversely, we see a reduction in PIP_2_ binding affinity with the K67N transformation (Fig. 3b and Supplementary Table 4). This quantitatively confirms that both the E179K and E179A mutations increase PIP_2_ channel affinity, whereas the K67N mutation decreases channel affinity. This agrees with the patient phenotypes: E179K causes NDM, i.e. an increase in channel activity, whereas K67N causes CHI, a reduction in channel activity.

**Fig. 3.**
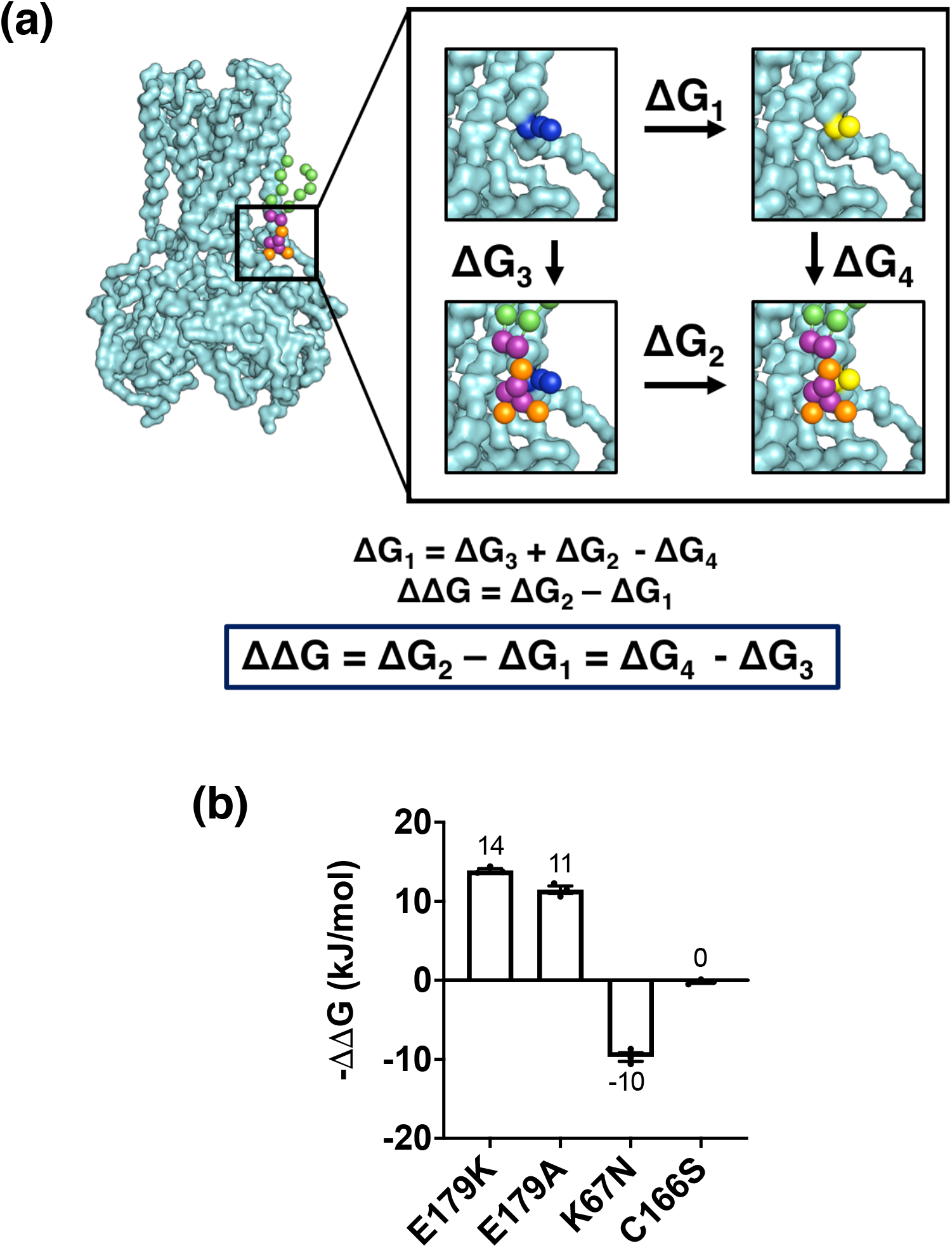
Free energy calculations using disease associated Kir6.2 mutations. **a** Schematic diagram showing the free energy calculation. An amino acid residue (blue sphere) is transformed into another residue (yellow sphere) between two states (PIP_2_ bound and free). **b** The energetic cost of making the residue mutation based on the schematic diagram (a).

As a control, we investigated an NDM mutation, C166S, which is distant from the PIP_2_ binding site. We expected that this mutation would have no impact on the PIP_2_ binding affinity, despite having an influence on the channel opening probability^29^. When perturbing the site using single residue FEP, we see effectively no change in PIP_2_ affinity of the channel, and therefore, unlike the E179A/K mutations, the C166S mutant does not appear to increase channel opening probability by increasing PIP_2_ affinity (Fig. 3c and Supplementary Table 2), but by a different mechanism.

### Experimental assessment of PIP_2_ binding to NDM mutant channels

We next assessed the neomycin sensitivity of the E179K mutant both in the presence and the absence of SUR1 to ascertain if the mutation increases both channel PIP_2_ affinity and channel activation (Fig. 4a-b). To enable expression of Kir6.2 without SUR1 we used a C-terminally truncated construct, Kir6.2ΔC36, which has been previously shown to express and traffic to the plasma membrane without SUR1^30^. Neomycin, a polycationic antibiotic, has previously been used as a tool to study Kir ion channel activation by PIP_2_ ^17^. While it does not bind to Kir channels directly, it acts by reversibly binding to PIP_2_, screening the charges and preventing binding^31^. A decrease in neomycin sensitivity (i.e. an increase in neomycin IC_50_) would indicate an increase in PIP_2_ dependent channel activation (i.e. PIP_2_ has a greater affinity for the Kir channel upon mutation). Another indication of an increase in channel activation by PIP_2_ is a slower rate of channel rundown and an increase in channel open probability^32^. We used these criteria to assess the effect of the SUR1 subunit on channel sensitivity to PIP_2_.

**Fig. 4.**
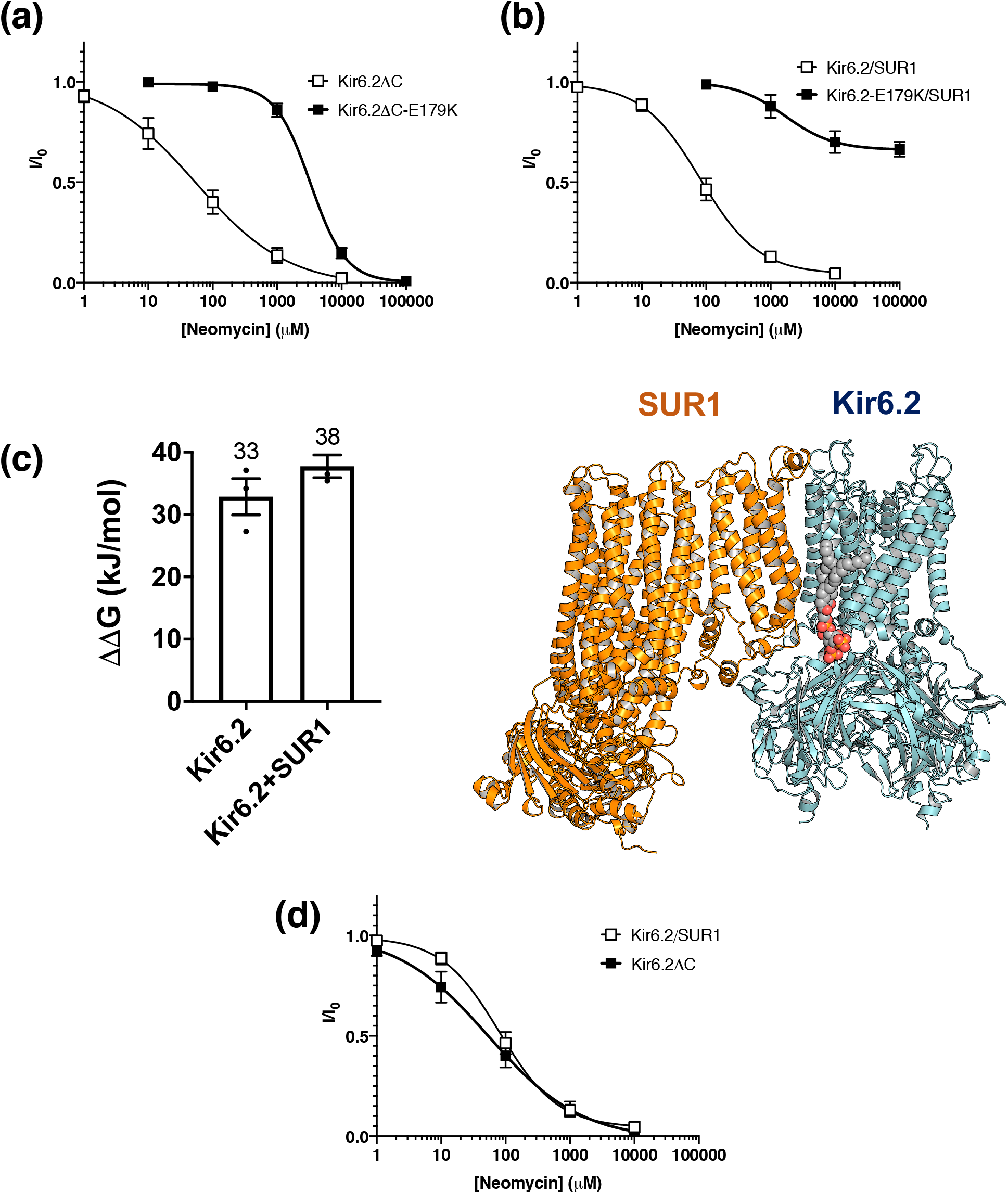
An effect of SUR1 subunit and NDM mutation on PIP_2_ affinity and activation. **a** Mean relationship between the neomycin concentration and the K_ATP_ current (I), expressed relative to the current in the absence of neomycin (I_0_) for Kir6.2ΔC (open squares, n=5) or hKir6.2ΔC-E179K channels (filled squares, n=5). **b** Mean relationship between the neomycin concentration and the K_ATP_ current (I) expressed relative to the current in the absence of neomycin (I_0_), for Kir6.2/SUR1 (open squares, n=5) or hKir6.2-E179K/SUR1 channels (filled squares, n=5). **c** (Left) Binding free energy between PIP_2_ and Kir6.2 ± SUR1. Values are rounded to the nearest whole number. Error bars represent the SEM (n=3) (Right) The PIP_2_ (grey) binding sites between hKir6.2 and SUR1. **d** Mean relationship between the neomycin concentration and the K_ATP_ current (I), expressed relative to the current in the absence of neomycin (I_0_), for Kir6.2 co-expressed with SUR1 (open squares, n=5) or hKir6.2ΔC expressed without SUR1 (filled squares, n=5).

We observed a 20-fold and a 50-fold increase in the IC_50_ value for neomycin block with the E179K mutation, both in the presence and absence of SUR1 respectively (Kir6.2-SUR1: IC_50_ = 81 μM, h=0.96, Kir6.2-E179K-SUR1: IC_50_ = 1.7 mM, h= 1.13, Kir6.2ΔC, IC_50_ = 55 μM, h= 0.61 and Kir6.2ΔC-E179K, IC_50_ = 3.3 mM, h = 1.6) This suggests an increase in the PIP_2_ sensitivity of the E179K variant channel both in the presence and in the absence of the SUR1 subunit, in agreement with our free energy calculations. Interestingly, in the presence of SUR1, channels with the Kir6.2-E179K mutation failed to fully close even in the presence of a very high neomycin concentration (0.1 M) (Fig. 4b). This suggests that the E179K mutation, in the presence of SUR1, may interfere with the channel gating mechanism independently of PIP_2_ action.

### Assessment of the PIP_2_ activity dependency on SUR1 subunit

Next, we experimentally assessed the importance of the SUR1 subunit for PIP_2_ binding affinity and activation. Previous studies suggested that the presence of SUR1 enhances the Po of Kir6.2ΔC^33,34^. However, the contribution of PIP_2_ to this modulation and the relationship between SUR1 and PIP_2_ sensitivity remains unclear. To address this issue, we calculated the relative binding free energy of hKir6.2 and PIP_2_ in both the presence and absence of the SUR1 subunit. Here, we show that addition of SUR1 only marginally increases the PIP_2_ binding free energy (Fig. 4c). Therefore, this result suggests that SUR1 has only a minor contribution to PIP_2_ affinity even though the SUR1 is only approximately 8 Å away from PIP_2_ headgroup.

To confirm that SUR1 has no effect on channel activation, we cloned and expressed Kir6.2 with SUR1 and Kir6.2ΔC in *Xenopus* oocytes and assessed the neomycin sensitivity of the channel. We found there was no significant difference in channel neomycin sensitivity in the presence and absence of SUR1 (Kir6.2/SUR1: IC_50_ = 81 μM, h = 0.96, Kir6.2ΔC, IC_50_ = 54 μM, h = 0.61) (Fig. 4d). This is in qualitative agreement with lack of a change in the free energy of PIP_2_ binding calculated from our FEP calculations.

### PIP_2_ binding affinity to other human inward rectifying potassium (hKir) channels

We next assessed the binding of PIP_2_ across other hKir channels by calculating the binding free energy of PIP_2_ perturbation to PC for the human Kir1.1, Kir2.2 and Kir3.2 (hKir1.1, hKir2.2 and hKir3.2) channels using the thermodynamic cycle described in Fig. 2a. The electrophysiological behaviour of these channels on PIP_2_ binding is well characterised^9^. We therefore perturbed PIP_2_ to PC, the dominant phospholipid species in the eukaryotic plasma membrane. Note that a potential of mean force (PMF) calculation shows the binding energy of PC to hKir6.2 is 0 ± 2 kJ/mol (Supplementary Fig. 9). Due to the absence of human Kir1.1 and human Kir2.2 structures in the PIP_2_ bound conformation, we generated hKir molecular models of both, as described in the Materials and Methods. The PIP_2_ binding sites and interacting residues on these proteins are all highly conserved (Fig. 5a-c, Supplementary Fig. 10).

**Fig. 5.**
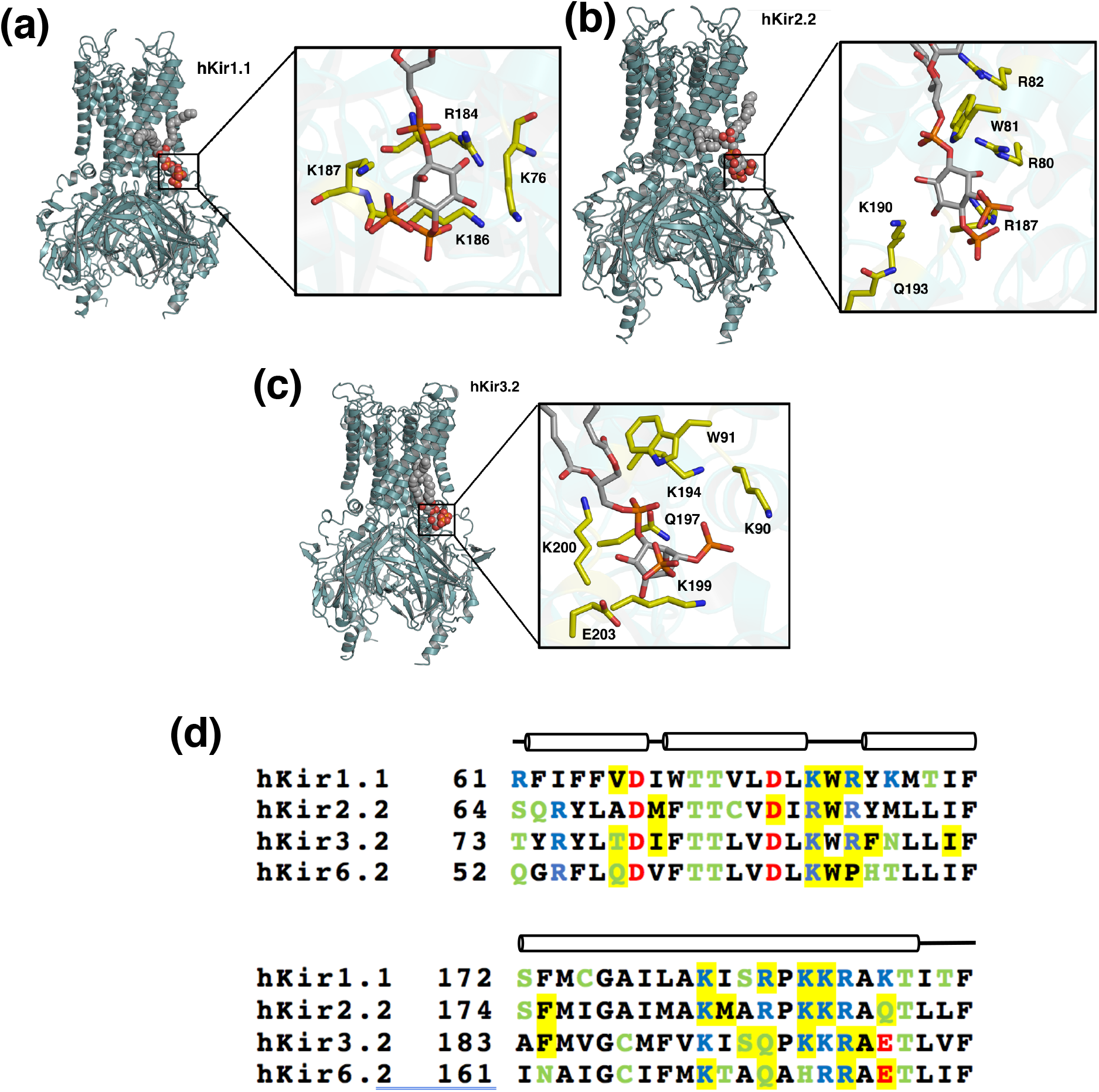
Free energy calculations for different hKir channels. Models of **a** hKir1.1, **b** hKir2.2 and **c** hKir3.2 channels in the PIP_2_ bound conformations after 1 μs of CG simulation and converted back to an atomistic description. Insets: Carbons of key PIP_2_ binding residues are highlighted in yellow, with PIP_2_ otherwise shown in CPK colours. **d** Sequence alignment between hKir1.1, hKir2.2, hKir3.2 and hKir6.2 channels on the region where the contacts are conserved between more than two channels. Highlighted in yellow are residues that contact PIP_2_ for more than 70% of the 1 μs simulations (n=5). Long cylinder represents an α-helix in the secondary structure and the line represents either disordered region or a kink within the α helix. Acid residues (Asp, Glu) are shown in red, basic residues (Lys, Arg) are shown in blue, Polar residues (Ser, Thr, Cys, Gln, Asn) are shown in green. Other residues are shown in black.

Our data reveal that the binding free energy of PIP_2_ to hKir1.1, hKir2.2 and hKir3.2 channels is higher than that to hKir6.2 channels (Fig. 6a and Supplementary Table 5). In our study, the PIP_2_ binding free energy value described here for hKir2.2 is similar to that previously recorded for the chicken Kir2.2 channel, which was reported as *ca.* −46 kJ/mol both using FEP and PMF^6,35^. The higher affinity of hKir3.2 was rather unexpected, as hKir3.2 has an equivalent glutamate in the same position at E179 on hKir6.2 (denoted as E201) (Fig. 5d). However, the minimum distance between the PIP_2_ headgroup and hKir3.2-E201 (*ca.* = 8.5 Å) is greater than that of than that of hKir6.2-E179 (*ca.* = 5 Å) in our molecular models.

**Fig. 6.**
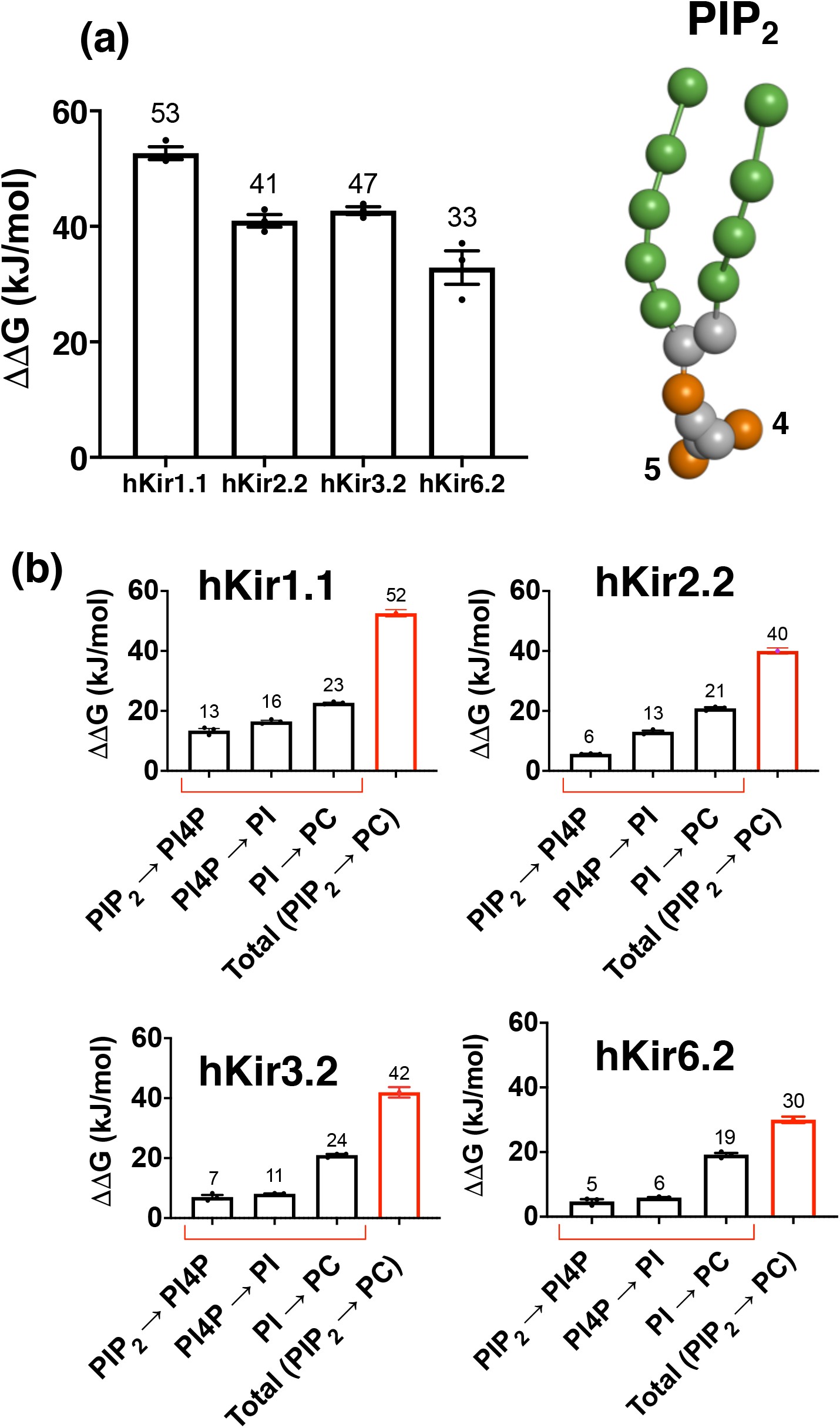
Free energy calculations for different hKir channels. **a** Binding free energy changes between different hKir channels as PIP_2_ is perturbed to PC. (n=3). **b** Binding free energy changes between PIP_2_ and hKir1.1, hKir2.2, hKir3.2 or hKir6.2 as each PIP_2_ phosphate group is sequentially perturbed: from PIP_2_ to PI4P, then to PI and finally to PC (black). The sum of the free energy change from PIP_2_ to PC is shown in red. Values are rounded to the nearest whole number. Error bars represent the SEM (n=3).

In order to account for the free energy difference between PIP_2_ and PC to all hKir channels, we examined the contribution of the individual phosphate and inositol groups of PIP_2_ in the binding to each of the channels, as described above (Fig. 2a). First, we show that the binding energy contributed by the 5’ phosphate is higher in hKir1.1 than the other three channels (PIP_2_>PI4P; Fig. 6b and Supplementary Table 6). In addition, we observed that 4’ phosphate and the inositol ring contribution is stronger in all other channels relative to hKir6.2 (PI4P>PI; Fig. 6b and Supplementary Table 6). Overall, this accounts for most of the energy differences as we perturbed PIP_2_ to PC in hKir1.1, hKir2.2 and hKir3.2. For both single- and multi-step approaches, hKir6.2 exhibited the lowest binding free energy for PIP_2_.

## Discussion

Here we build on our recent application of the CG-FEP approach for comparing the binding free energy between two lipid species to a given site on a membrane protein^6^. We show that this approach enables us to complete a thermodynamic cycle where the sum of the individual perturbation steps (i.e. PIP_2_>PI4P>PI>PC) is equivalent to the single-step transformation from PIP_2_ to PC (Supplementary Fig. 6). The results of our application of the free-energy calculations are within the range of values that are commonly observed for PIP_2_ interactions with membrane proteins^6,35–37^. One of our concerns is the ability of the CG forcefield to distinguish between similar inositol lipids in the free energy calculations. This application of CG-FEP has demonstrated that the method can show differences between PIP_2_, PI4P, PI and PC at the accuracy of at least 5 kJ/mol (~1.5 k_B_T). This complements the previous free energy calculations, which show that the CG forcefield is able to distinguish between PIP_2_ and PIP_3_^37^. Overall, this illustrates the robustness of the application and demonstrates its power as a relatively cheap and effective *in silico* approach for comparing lipid-binding free energies to a membrane protein of interest. Nevertheless, although there is good agreement between the single- and multi-step approaches we cannot exclude the possibility that the individual particle contributions may be either over- or under-estimated due to the CG approach.

In addition, we demonstrate that amino acid mutations can also be investigated using CG-FEP, thus allowing us to probe the effect of a given amino acid’s substitution on lipid binding. This is an extension to the traditional atomistic FEP mutation approach^21^, which enables convergence to be achieved more quickly and easily^21^. It also allows binding to be much more easily measured than in *vitro* lipid binding studies. Thus, the application of CG-FEP is potentially a valuable tool for the relatively high-throughput analysis of multiple disease-causing mutations that are related to lipid binding.

To test the capabilities of these methodologies, we applied them to the K_ATP_ channel (Kir6.2/SUR1), a biologically important potassium channel that is implicated in insulin secretion. We used a combination of the above methods to demonstrate that clinically identified mutations in the Kir6.2 subunit (causing NDM and CHI) affect the affinity of the channel for different PIP lipids. In addition, we demonstrated that the neonatal diabetes mutations - E179K and E179A - which lie near the PIP_2_ binding site result in channel gain of function by enhancing PIP_2_ binding (and Po). We present both computational and electrophysiological data which are in good agreement, thereby demonstrating that this application of an existing method provides a potentially powerful method for the scanning and annotating of how disease-associated mutations modulate lipid binding to channels.

## Methods

### Molecular modelling

Modeller9v16^38^ was used to add the missing loops and amino acid residues to the cryo-EM structure of the Kir6.2 channel [PDB entry: 6BAA]^39^ and to generate a human Kir6.2 model based on residues 32-352. In both the cryo-EM structure and our model there are 32 amino acids missing at the N-terminus and 39 at the C-terminus. Modeller9v16 was also used to generate a model of the human K_ATP_ channel octamer (hKir6.2 tetramer + four SUR1 [PDB entry: 6BAA]) and the models of the hKir6.2 mutant channels. Each model was compared to its initial template structure to ensure that the modelling had not demonstrably altered the original secondary structure or the rotation of the amino acid sidechains (overall RMSD of all protein atoms < 1.0 Å). Models of the other hKir channels were generated using Swiss-Model^40^, with human Kir1.1 and Kir2.2 based on 3SPH and human Kir3.2 based on 3SYA. The chicken Kir2.2 structure with bound diC8-PIP_2_ [PDB entry: 3SPI]^12^ was used to dock PIP_2_ to hKir6.2.

### Coarse-grained (CG) Simulations

All protein structures were converted to their CG representation and embedded in a PC (1-palmitoyl-e-oleoyl-sn-glycero-3-phosphocholine) bilayer using the self-assembly MemProtMD protocol^24,41^ and the MARTINI v2 biomolecular forcefield^4^. This approach orients the structure of the transmembrane protein parallel to the z-axis using MEMEMBED^42^. The protein is then placed in a periodic box at minimum distance of 30 Å from the edge of the box in both x and y directions, and with a z dimension of 80 Å. The structure is then converted to a CG representation with *martinize.py* with an application of an elastic network with a force constant of 1,000 kJ/mol/nm^2^ between backbone beads within 0.5-0.9 nm to maintain their secondary and tertiary structure. The PC lipid is then added to the periodic box, allowing them to assemble freely around the protein. The z-dimension of the box is then extended so that the minimum distance between the protein and the face of the box is 30 Å apart, and then flooded with the coarse-grain water particles, Na^+^ and Cl^−^ ions to a final concentration of 0.15 M to neutralize the system. The total number of the molecules in the setup is described in the supplementary table 8. A temperature of 323 K was maintained with V-rescale temperature coupling^43^, while 1 atm pressure was controlled using semi-isotropic Parrinello-Rahman pressure coupling^44^. Systems were energy minimised using the steepest descents algorithm and equilibrated for 5 ns with 1,000 kJ/mol/nm^2^ position restraints on backbone beads, prior to 1 μs production. All simulations, root mean square deviation (RMSD) calculations and distance analyses were carried out using GROMACS v2018^45^ and all structural alignments and docking were carried out using PyMOL.^46^

### Free energy perturbation (FEP) calculation of PIP lipids

The hKir6.2 tetramer with one bound PIP_2_, obtained after equilibration, was used as the initial co-ordinates for the majority of the FEP calculations. Here, we calculate a relative binding free energy (ΔΔG) by converting from one lipid type (such as PIP_2_) to a series of other phospholipids (such as PIP_2_, PI4P, PI and PC) along a reaction coordinate in a chemical space denoted λ (Fig. 2a). As is standard for FEP calculations, separate transformations were performed with either the lipid bound to the channel or in bulk membrane.

We applied FEP to hKir6.2, hKir3.2, hKir2.2 and hKir1.1 and the following pairs of inositol lipids: (PIP_2_ and PI4P), (PI4P and PI), (PI and PC) and (PIP_2_ and PC). This allows us to create a thermodynamic cycle for the different lipids of interest (Supplementary Fig. 6). For these, specific phosphate and inositol sugar particles were transformed into a dummy particle with no interaction properties, in a stepwise process as described in Fig. 2b. Coulombic (charge interactions) and Lennard-Jones (van der Waals interactions) were turned off separately, with a soft-core parameter used for the Lennard-Jones interactions. The coulombic interactions were perturbed linearly (λ = 0, 0.1, 0.2 … 0.9, 1.0) in the first 10 simulation windows, with the van der Waals interactions perturbed linearly (λ = 0, 0.1, 0.2 …… 0.9, 1.0) in the last 10 simulation windows with the soft-core parameters of α = 0.5 and σ = 0.3. Each simulation window was energy minimised and equilibrated as described above, before three production runs were carried out for 250 ns with randomised initial velocities, using a leap-frog stochastic dynamics integrator. A flat-bottom distance restraint between the PO4 phosphate group and protein backbone beads at 6 Å radii from the PIP molecule was applied using Plumed (1000 kJ/mol/nm^2^, 8 Å cut-off)^27^. This prevents the bound lipid from drifting away from its binding pocket and increases the accuracy of the calculation (Supplementary Fig. 1b,1c). The free energy pathways were constructed using the *alchemical-analysis* software package^47^, where the energies are calculated based on the 300 ns of the data for a good convergence^6^. Thus, a total of 642.6 μs simulations were performed for the FEP calculations performed in this study. Analyses was run using the Multistate Bennett Acceptance Ratio (MBAR)^25^. All values are reported are reported as mean ± SEM. All simulations were carried out using GROMACS v2018^45^.

### Free energy perturbation (FEP) between amino acid residues

As before, the PIP_2_-bound equilibrated hKir6.2 system was used as the initial co-ordinates for the FEP calculations. When assessing the influence of ND or CHI mutations, we calculated changes in the relative binding free energy (ΔΔG) by alchemically transforming the wild-type amino acid residue to its mutant counterpart. This was performed using a change in chemical space denoted as λ, and applying the previously described protocol. The series of transformations carried out are shown in Supplementary Table 7. Additional simulations were run by performing the same perturbation of the protein in a bulk POPC membrane in the absence of PIP_2_. The ΔΔG terms were then calculated as described in Fig. 3b.

### Potential of Mean Force Calculation (PMF)

PMF calculations were set up similarly to that described previously^6^. The protein was built in the POPC bilayer, and a single POPC lipid was pulled from the binding site using steered-MD, where the collective variables (CV) are the distance between the lipid headgroup and the centre of mass of the protein. The initial position for POPC was modified from an initial PIP_2_ co-ordinate. The simulations were calculated along the CV at 0.2 Å interval for optimal histogram overlap, with a 1000 kJ/mol/nm^2^ umbrella potential applied to restrain the position of the lipid along the CV. Positional restraints of 100 kJ/mol/nm^2^ were applied to the protein backbone to prevent rotation of the protein in the bilayer. For each window, the simulations were run for 500 ns, which was sufficient to see convergence. Thus, this adds up to a total of 20 μs for the PMF calculations. The 1D energy profile was generated using weighted histogram analysis method (WHAM) using the *gmx wham* tool (200 rounds Bayesian Bootstrap) ^48^.

### Molecular biology

Human Kir6.2 (Genbank NM000525) and human SUR1 (Genbank NM_000352.5) were cloned into the pBF vector. Site-directed mutagenesis was performed using QuickChange XL (Stratagene), followed by synthesis of capped mRNA using mMESSAGE (Invitrogen). All constructs were validated by restriction digest and DNA sequencing (MRC I PPU, School of Life Science, University of Dundee, Scotland). *Xenopus laevis* oocytes were prepared as previously reported ^49^. The oocytes were co-injected with ∼4 ng of SUR1 mRNA and ∼0.8 ng wild-type or mutant Kir6.2 mRNA. In some experiments, oocytes were injected with wild-type or mutant Kir6.2 possessing a C-terminal 36 amino acid truncation (Kir6.2ΔC) mRNA, which allows surface membrane expression^33^. Oocytes were incubated in Barth’s solution and studied 1–4 days after injection.

### Electrophysiology

Inside-out patch-clamp recordings were performed using an EPC7 amplifier (List Electronik) at a constant holding potential of −60 mV. The pipette solution contained 140 mM KCl, 1.2 mM MgCl_2_, 2.6 mM CaCl_2_, 10 mM HEPES (pH 7.4 with KOH). For experiments with neomycin, the Mg-free intracellular solution contained 107 mM KCl, 10 mM HEPES and 10 mM EDTA (pH 7.2 with KOH). To account for possible rundown, the control current (*I*_C_) was taken as the mean of the current in control solution before and after neomycin application. Concentration– response curves were fitted with a modified Hill equation (1):

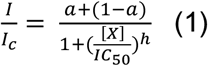

where [*X*] is the concentration of the test substance, IC_50_ is the concentration at which inhibition is half maximal, *h* is the slope factor (Hill coefficient) and *a* represents the fraction of unblocked current at saturating neomycin. Single-channel currents were recorded at −60 mV, filtered at 5 kHz, sampled at 20–50 kHz, and analysed using a combination of Clampfit (Axon Instruments) and GraphPad Prism 8. Data are given as mean ± SEM.

## Acknowledgements

We thank Mark Sansom for fruitful discussions and Irfan Alibay, Josh Sauer, and Owen Vickery for advice and technical support. TP holds a Wellcome Trust OXION studentship and a Clarendon scholarship. Research in PJS’s lab is funded by Wellcome (208361/Z/17/Z), the MRC (MR/S009213/1) and BBSRC (BB/P01948X/1, BB/R002517/1 and BB/S003339/1). Research in FMA’s lab is funded by the MRC (MR/T002107/1), BBSRC (BB/R002517/1, BB/R017220/1) and Wellcome Trust (102161/Z/13/Z). This project made use of time on ARCHER and JADE granted via the UK High-End Computing Consortium for Biomolecular Simulation, HECBioSim (http://hecbiosim.ac.uk), supported by EPSRC (grant no. EP/R029407/1). PJS acknowledges Athena at HPC Midlands+, which was funded by the EPSRC on grant EP/P020232/1, and the University of Warwick Scientific Computing Research Technology Platform for computational access.

## Author contributions

T.P. prepared the mutants for the electrophysiological studies and performed the coarse-grained molecular dynamics simulations and free energy calculations. P.P. provided recording from the electrophysiology experiments. T.P., R.A.C., F.M.A and P.J.S. jointly designed the experiments, analysed the data and wrote the manuscript.

## Data availability

All data are available from the corresponding author on request.

## Supplementary information

**Supplementary Figure 1.**
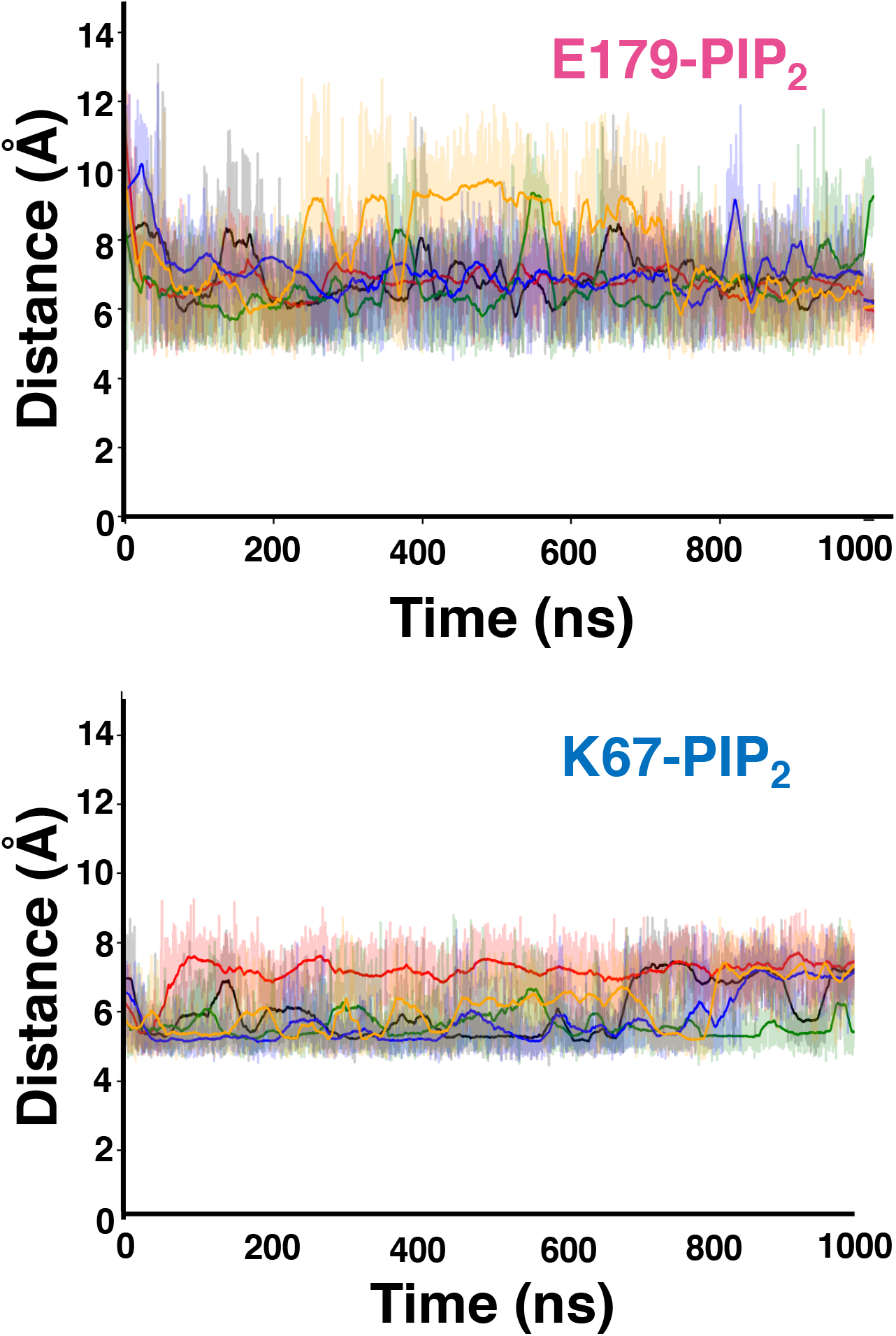
Distance between protein backbone and the PIP_2_ headgroup. Calculated distance between the PIP_2_ headgroup and the backbone of either **a** E179 or **b** K67, over a 1 μs simulation. The different colours indicate the individual repeats of the simulation. The darker lines show the running average for each simulation (n=5).

**Supplementary Figure 2.**
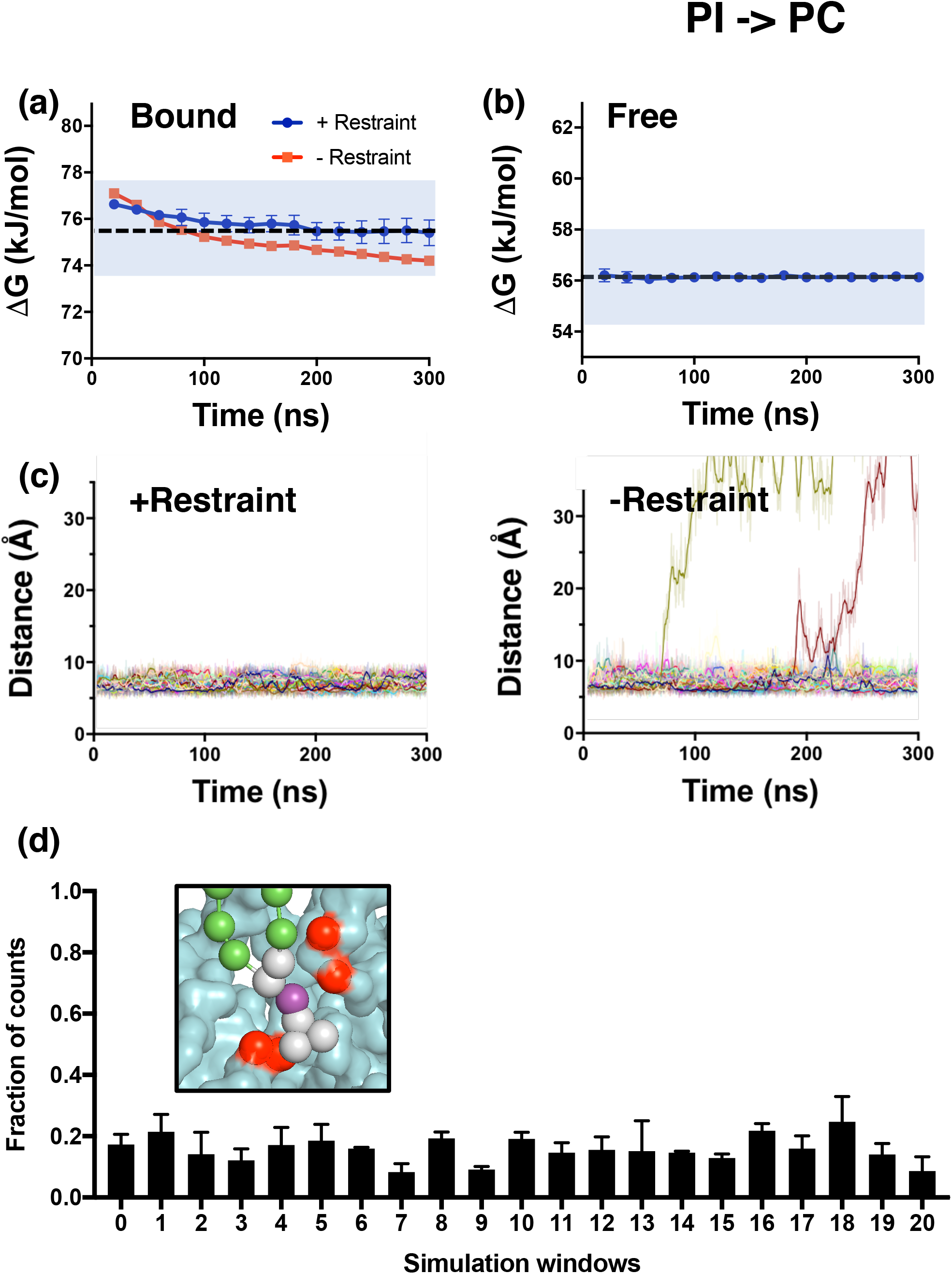
Convergence analysis of PI to PC free energy perturbation. **a** Convergence of the free energy calculation during PI to PC perturbation in the presence of hKir6.2. The analysis was carried out with (blue) and without (red) the flat-bottom restraint. The blue shaded region illustrates the thermal fluctuation of the system, i.e. kT. **b** Convergence of the free energy calculation during PI to PC perturbation in the bulk PC bilayer in the absence of hKir6.2. The blue shaded region illustrates the thermal fluctuation of the system, i.e. kT. **c** The distance between the phosphate headgroup (PO4 particle) and the average position of four amino acid residues (red), in the presence and absence of a flat-bottom restraint. These four residues were chosen as they are 6 Å away from the lipid headgroup. Different colours represent the simulations in the different alchemical states (λ windows) of PI to PC transformation. **d** Fraction of counts where the PO4 particle experiences the flat-bottom restraint in each simulation window. Inset: A flat bottom restraint was imposed between the PO4 particle (purple) and the protein backbone (red).

**Supplementary Figure 3.**
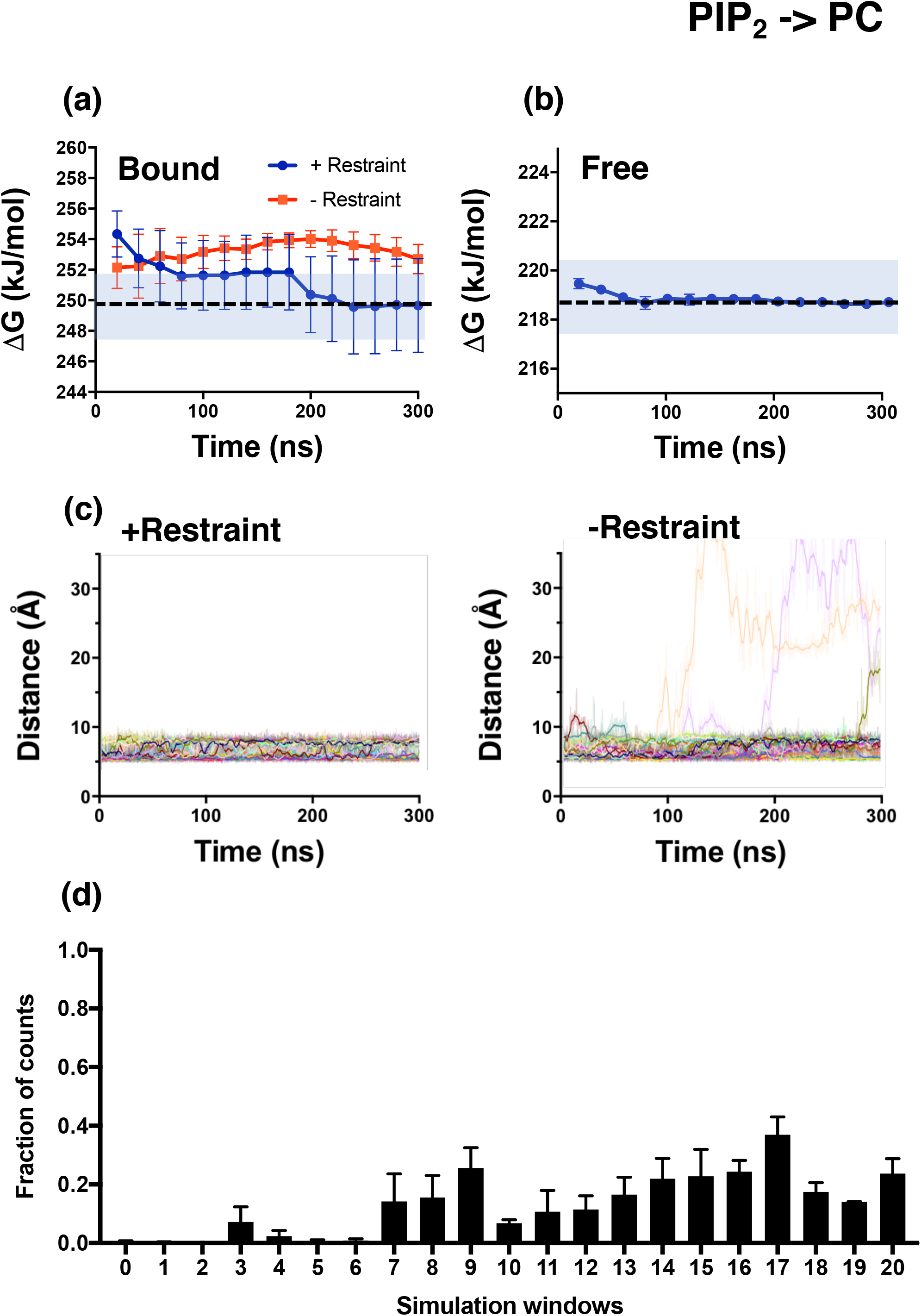
Convergence analysis of PIP_2_ to PC free energy perturbation. **a** Convergence of the free energy calculation during PIP_2_ to PC perturbation in the presence of hKir6.2. The analysis was carried out with (blue) and without (orange) the flat-bottom restraint. The blue shaded region illustrates the thermal fluctuation of the system, i.e. kT. **b** Convergence of the free energy calculation during PIP_2_ to PC perturbation in the bulk PC bilayer. The blue shaded region illustrates the thermal fluctuation of the system, i.e. kT. **c** The distance between the phosphate headgroup (PO4 particle) and the centre position between the backbone of 4 amino acid residues (red) in the presence and absence of a flat-bottom restraint. These residues were chosen as they are 6 Å away from the lipid headgroup. Different colours represent the simulations in the different alchemical states (λ windows) of PIP_2_ to PC transformation. **d** Fraction of counts where the PO4 particle experiences the flat-bottom restraint in each simulation windows.

**Supplementary Figure 4.**
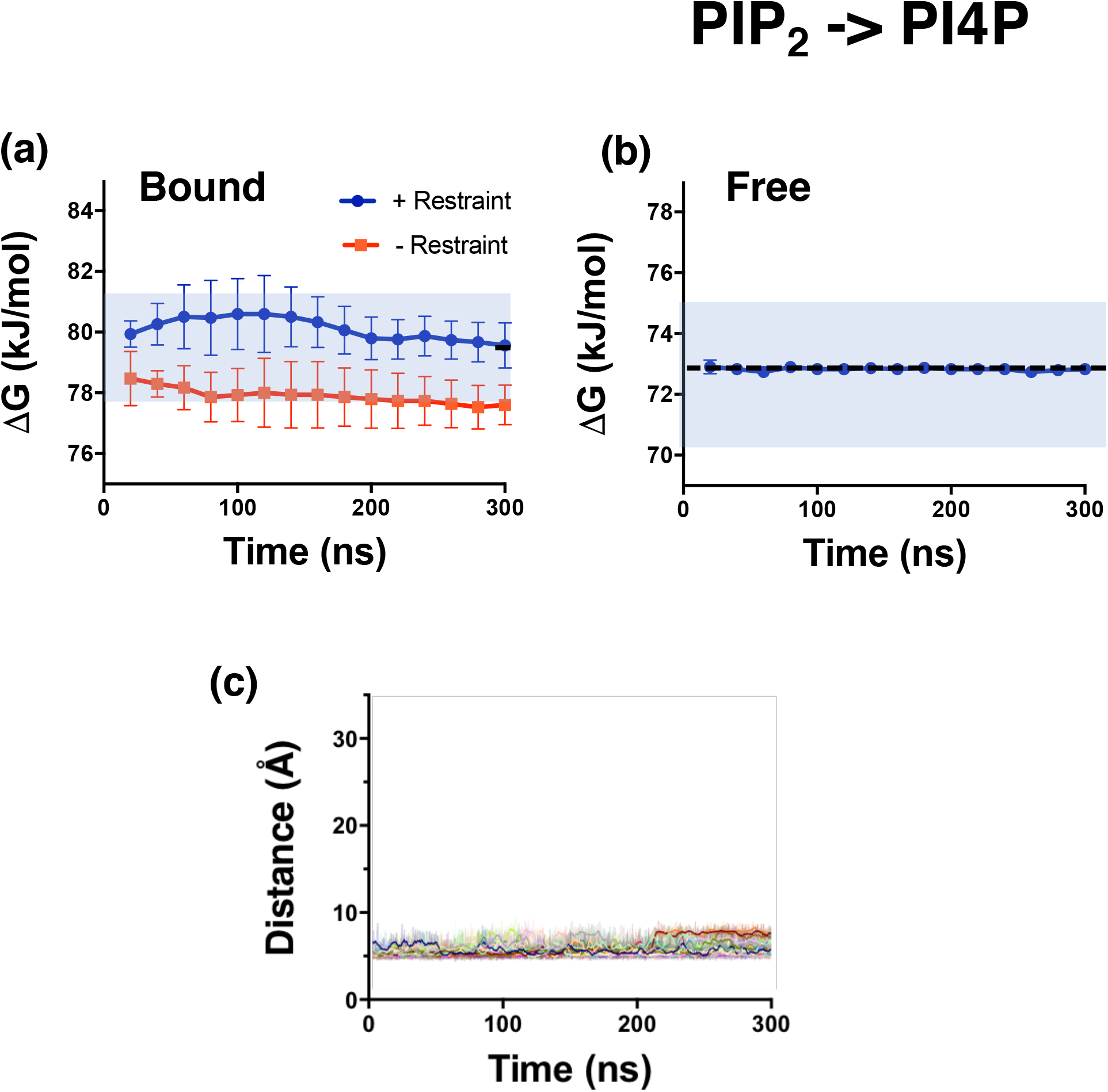
Convergence analysis of PIP_2_ to PI4P free energy perturbation. **a** Convergence of the free energy calculation during PIP_2_ to PI4P perturbation in the presence of hKir6.2. The analysis was carried out with (blue) and without (orange) the flat-bottom restraint. The blue shaded region illustrates the thermal fluctuation of the system, i.e. kT. **b** Convergence of the free energy calculation during PIP_2_ to PI4P perturbation in the bulk PC bilayer. The blue shaded region illustrates the thermal fluctuation of the system, i.e. kT. **c** The distance between the phosphate headgroup (PO4 particle) and the flat-bottom restraint. Different colours represent the individual simulations in the different alchemical states (λ windows) of the PIP_2_ to PI4P transformation.

**Supplementary Figure 5.**
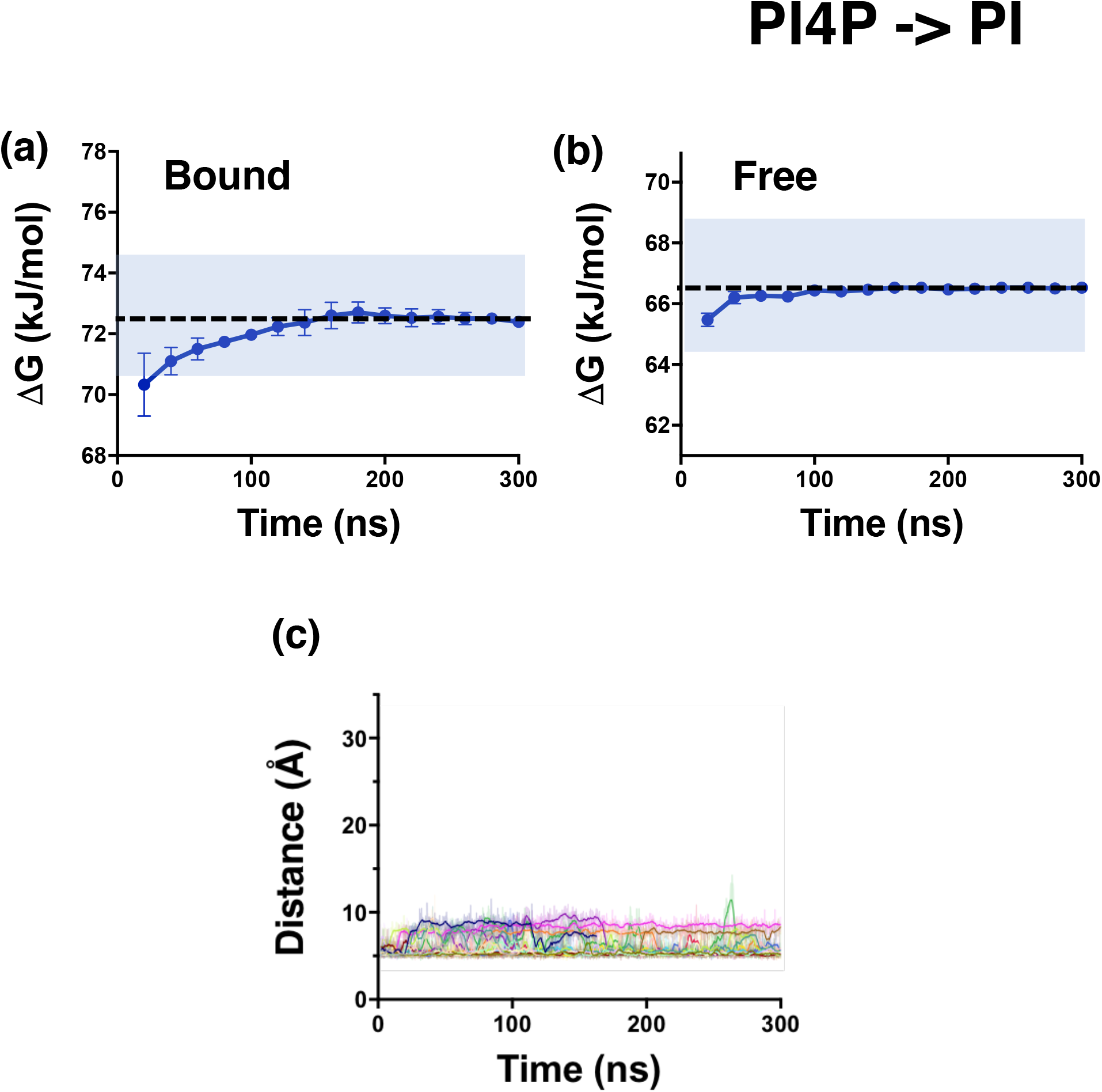
Convergence analysis of PI4P to PI free energy perturbation. **a** Convergence of the free energy calculation during PI4P to PI perturbation in the presence of hKir6.2. The analysis was carried out with (blue) and without (orange) the flat-bottom restraint. The blue shaded region illustrates the thermal fluctuation of the system, i.e. kT. **b** Convergence of the free energy calculation during PI4P to PI perturbation in the bulk PC bilayer. The blue shaded region illustrates the thermal fluctuation of the system, i.e. kT. **c** The distance between the phosphate headgroup (PO4 particle) and the flat-bottom restraint. Different colours represent the simulations in the distinct alchemical states (λ windows) of the PI4P to PI transformation.

**Supplementary Figure 6.**
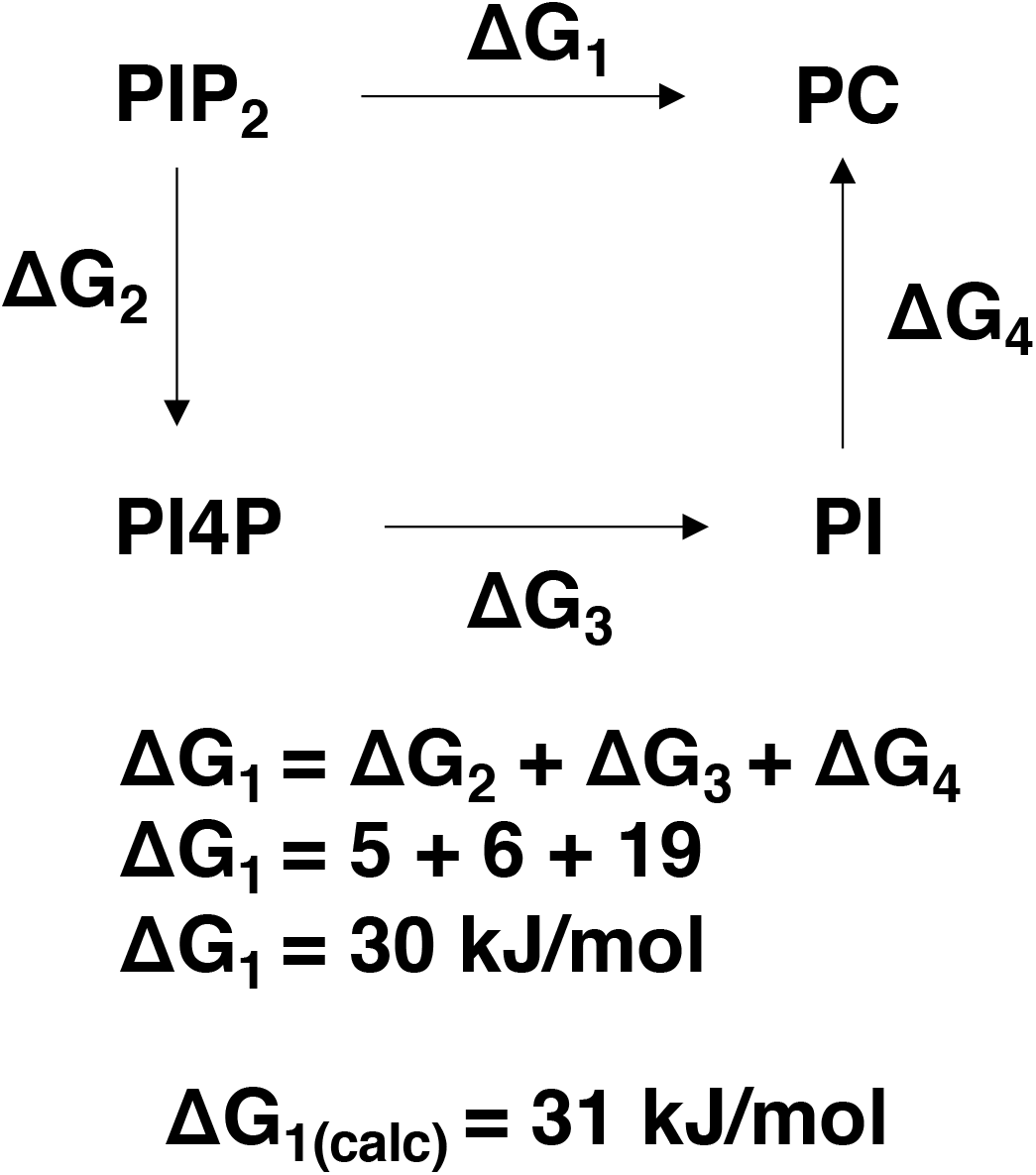
A complete thermodynamic cycle in PIP_2_ stepwise perturbation. The thermodynamic cycle representing relative binding free energy between phosphoinositide lipids (PIP_2_, PI4P and PI) and PC.

**Supplementary Figure 7.**
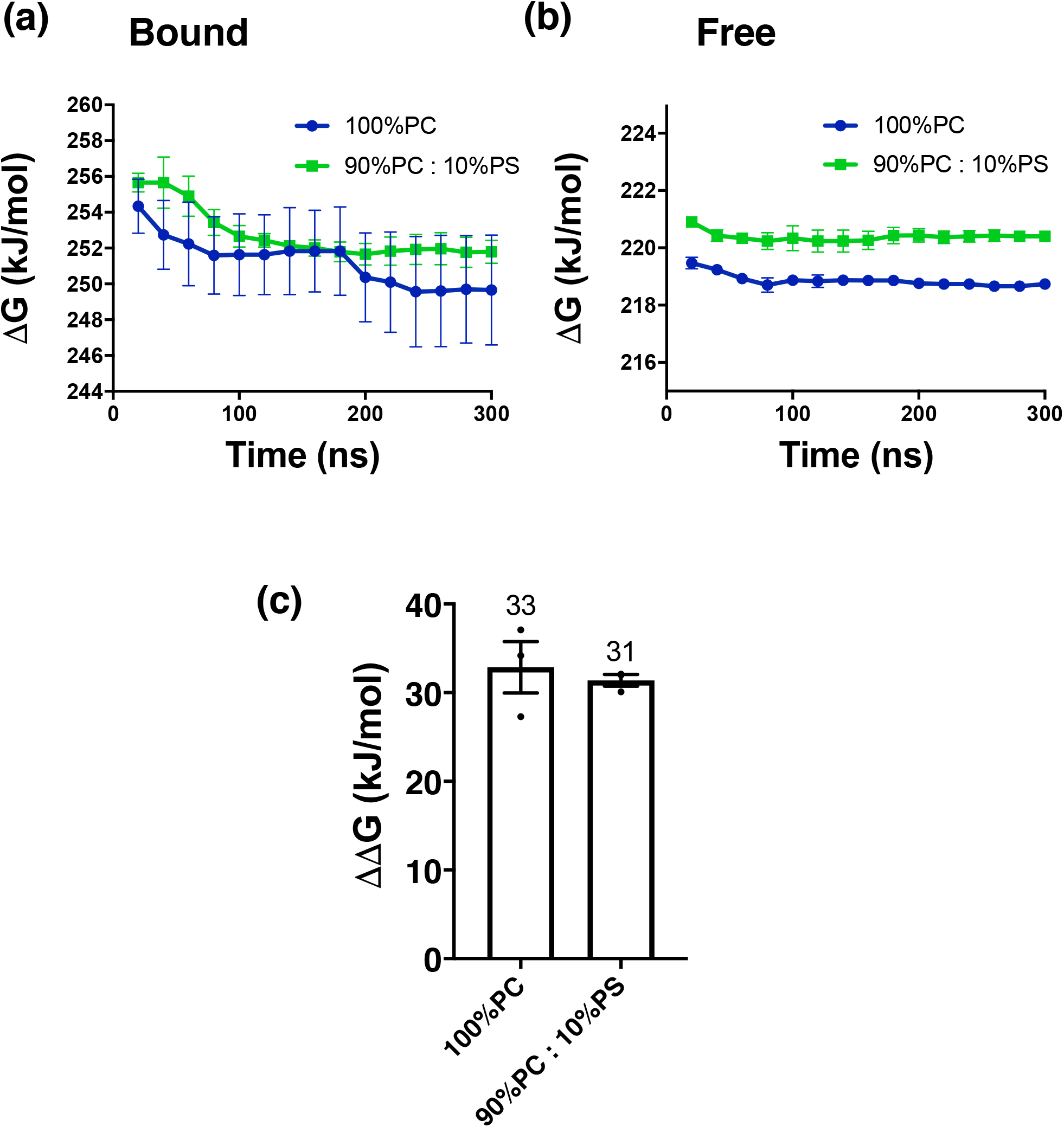
Convergence analysis of PIP_2_ to PC free energy perturbation in anionic lipid environment. **a** Convergence of the free energy calculation during PIP_2_ to PC perturbation in the presence of hKir6.2. The analysis was carried out in 100% PC bilayer (blue) or mixed bilayer containing 10% PS (green) **b** Convergence of the free energy calculation during PIP_2_ to PC perturbation in the bulk PC bilayer. The analysis was carried out in 100% PC bilayer (blue) or mixed bilayer containing 10% PS (green) **c** Binding free energy between PIP_2_ and Kir6.2 in 100% PC bilayer or in 10% PS. Values are rounded to the nearest whole number. Error bars represent the SEM (n=3)

**Supplementary Figure 8.**
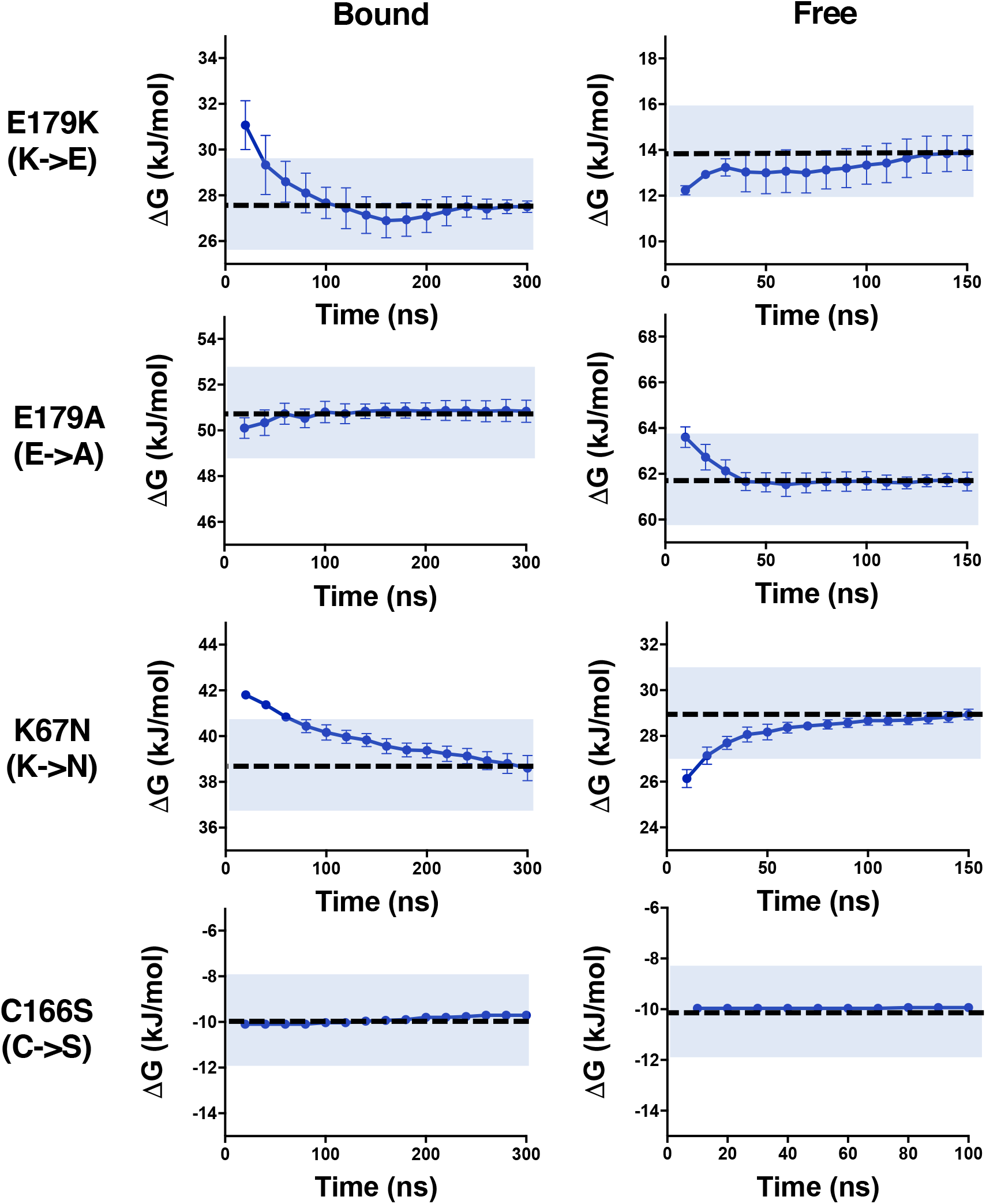
Convergence analysis of E179K (K→E), E179A, K67N and C166S free energy perturbation. Convergence of the free energy calculation during lysine to glutamate perturbation for residue 179 (E179K), glutamate to alanine perturbation for residue 179 (E179A), lysine to asparagine perturbation for residue 67 (K67N) and cysteine to serine perturbation for position 166 (C166S) of hKir6.2 in (a) the presence and (b) the absence of PIP_2_. The shaded blue region illustrates the thermal fluctuation of the system, i.e. kT.

**Supplementary Figure 9.**
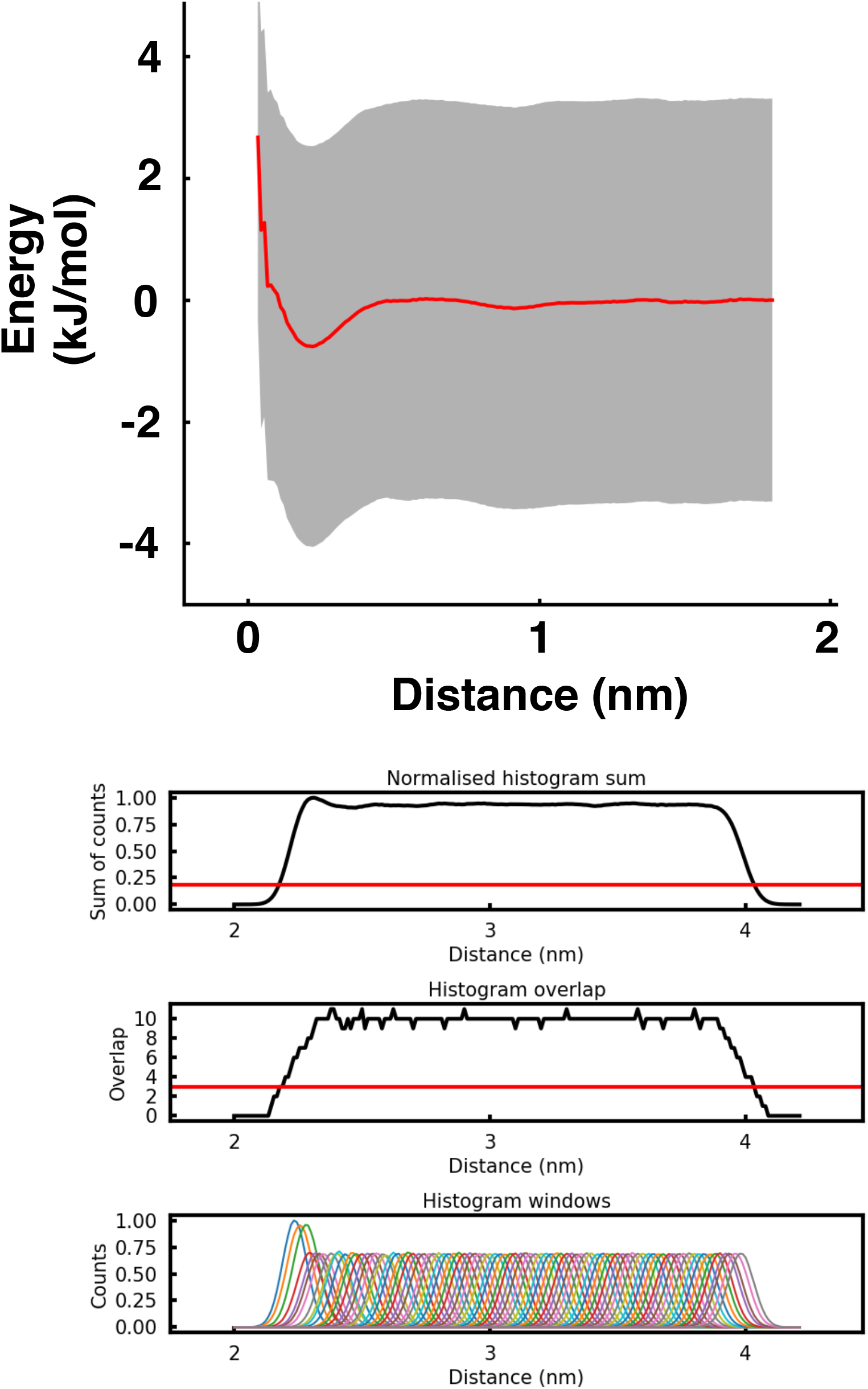
PMF data for PC binding to hKir6.2. The 1D energy landscape for PIP_2_ binding to hKir6.2. The y-axis is set to 0 in PC and the x-axis is set to 0 nm. The calculated ΔG is ca. 0 kJ/mol.

**Supplementary Figure 10.**
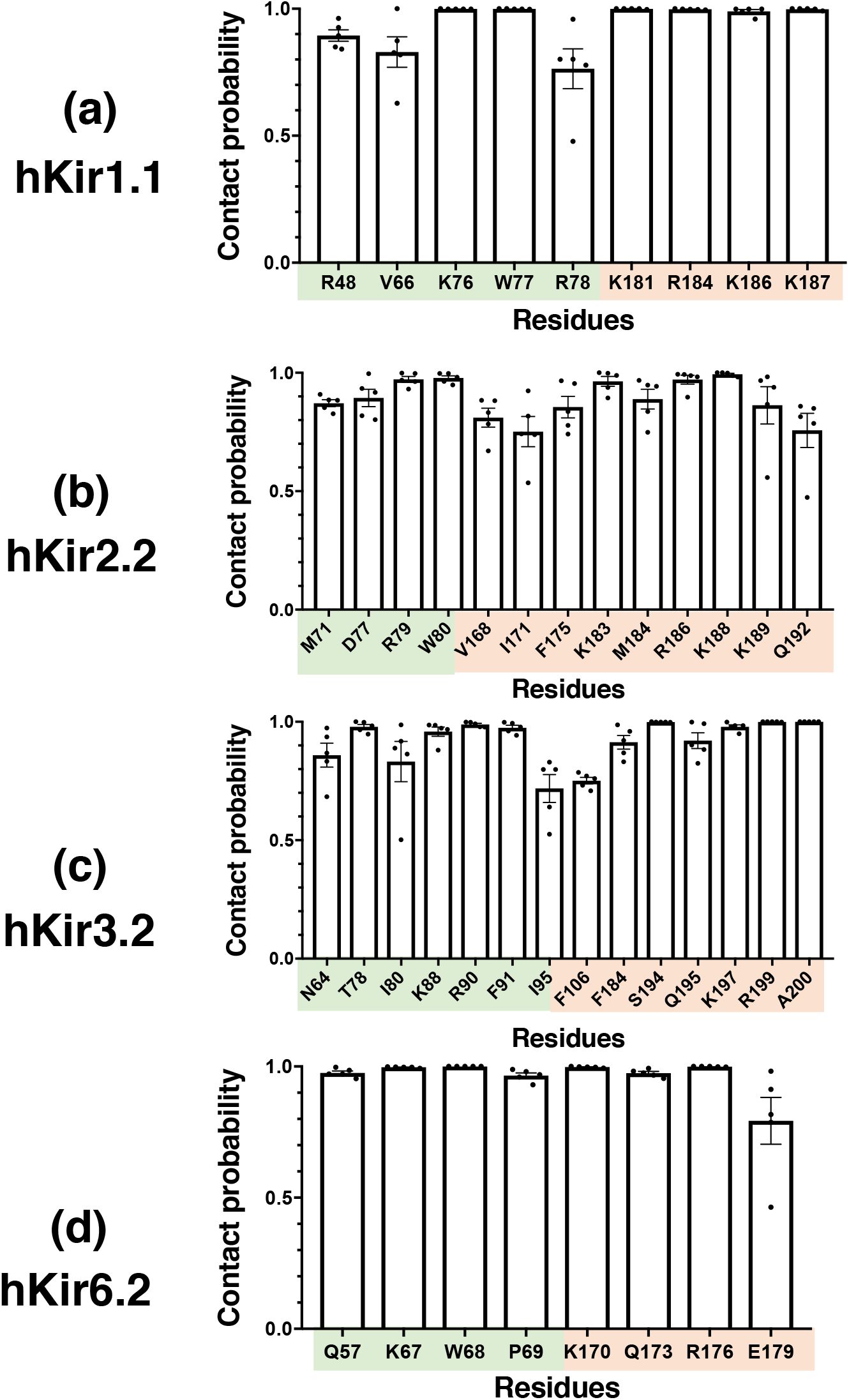
A PIP_2_ binding site on hKir channels. PIP_2_ contact analysis showing residues which make contact >75% of the time during a 1 μs simulation for **a** hKir1.1, **b** hKir2.2, **c** hKir3.2 or **d** hKir6.2 channels (n = 5). Residues in the N-terminal domain are highlighted in green and in the C-terminal domain are coloured in orange.

### Supplementary Tables

**Supplementary Table 1.**
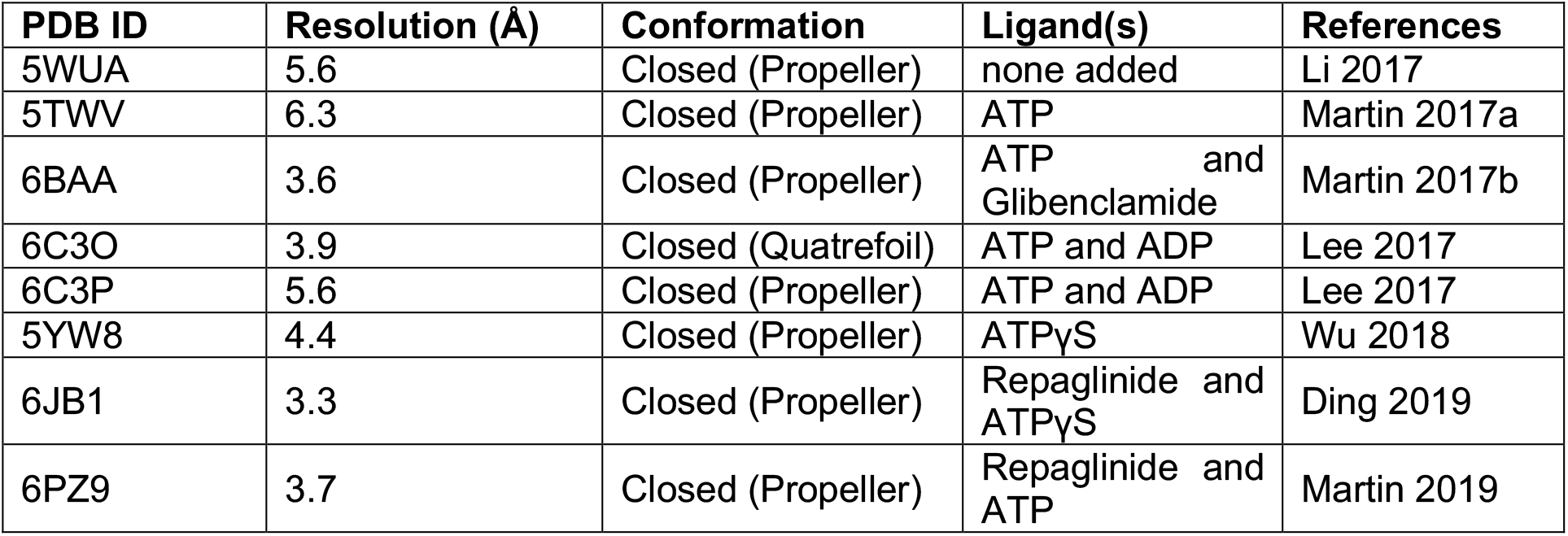
List of all cryo-EM K_ATP_ channel structures. Table of K_ATP_ channel structures giving their PDB ID, overall resolution, conformation and bound ligands.

**Supplementary Table 2.**
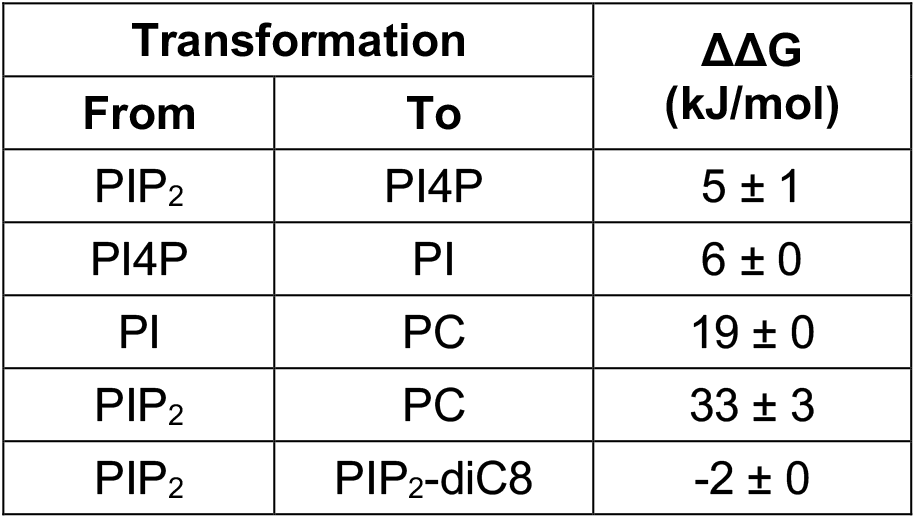
Calculation of the ΔΔG on an individual phosphate group and the fatty acid chains on a hKir6.2 tetramer. Change in binding free energy (ΔΔG) when individual phosphate groups are perturbed (i.e. from PIP_2_ to PI4P, from PI4P to PI and from PI to PC (values in black), when PIP_2_ is perturbed to PC and when PIP_2_ is perturbed to PIP_2_-diC8. Values are rounded to the nearest whole number (n=3).

**Supplementary Table 3.** Free energy calculations using PIP_2_ to PC transformation. Binding free energy changes between wild-type and mutated hKir6.2 channels and hKir6.2 channel with SUR1 as we perturbed PIP_2_ to PC. Values are rounded to the nearest whole number. Mean and SEM (n=3).

| Protein | $\Delta\Delta G$<br>(kJ/mol) |
| --- | --- |
| hKir6.2 | 33 ± 3 |
| hKir6.2 + SUR1 | 38 ± 2 |

**Supplementary Table 4.** Free energy calculations using disease associated hKir6.2 mutations. The energetic cost of making the residue mutation based on the schematic diagram (Figure 3b).

| **Mutation** | **-$\Delta\Delta G$**<br>**(kJ/mol)** |
| --- | --- |
| E179K (K→E) | 14 ± 1 |
| E179A | 11 ± 1 |
| K67N | -10 ± 1 |
| C166S | 0 ± 0 |

**Supplementary Table 5.** Free energy calculation on different hKir channels. Binding free energy changes between different hKir channels as we perturbed PIP_2_ to PC. Values are rounded to the nearest whole number. Mean and SEM (n=3).

| Protein | $\Delta\Delta G$<br>(kJ/mol) |
| --- | --- |
| hKir1.1 | 53 ± 1 |
| hKir2.2 | 41 ± 1 |
| hKir3.2 | 47 ± 1 |
| hKir6.2 | 33 ± 3 |

**Supplementary Table 6.** Calculation of the ΔΔG on an individual phosphate group on hKir channels. Change in binding free energy (ΔΔG) when individual phosphate groups are perturbed (i.e. from PIP_2_ to PI4P, from PI4P to PI and from PI to PC (values in black), The sum of these values are shown in bold. Values are rounded to the nearest whole number (n=3).

| Protein | Transformation | | $\Delta\Delta G$<br>(kJ/mol) |
| --- | --- | --- | --- |
|  | From | To |  |
| hKir1.1 | PIP<sub>2</sub> | PI4P | 13 ± 1 |
|  | PI4P | PI | 16 ± 0 |
|  | PI | PC | 23 ± 0 |
|  | Sum |  | 52 ± 1 |
| hKir2.2 | PIP<sub>2</sub> | PI4P | 6 ± 0 |
|  | PI4P | PI | 13 ± 0 |
|  | PI | PC | 21 ± 1 |
|  | Sum |  | 40 ± 1 |
| hKir3.2 | PIP<sub>2</sub> | PI4P | 7 ± 0 |
|  | PI4P | PI | 11 ± 1 |
|  | PI | PC | 24 ± 0 |
|  | Sum |  | 42 ± 1 |
| hKir6.2 | PIP<sub>2</sub> | PI4P | 5 ± 1 |
|  | PI4P | PI | 6 ± 0 |
|  | PI | PC | 19 ± 0 |
|  | Sum |  | 30 ± 1 |

**Supplementary Table 7.** Direction of free energy perturbation of amino acid residues. The direction of the transformation and the particle types for free energy perturbation analysis. The particles are suggested based on MARTINI2.2 forcefield. Dum is a dummy particle with a mass of 72 and no bonded or non-bonded interactions.

| Mutation | Direction of Perturbation | From |  | To |  |
| --- | --- | --- | --- | --- | --- |
|  |  | SC1 | SC2 | SC1 | SC2 |
| K67N | WT → K67N | C3 | Qd | P5 | Dum |
| C166S | WT → C166S | C5 | - | P1 | - |
| E179K | E179K → WT | C3 | Qd | Qa | Dum |
| E179A | WT → E179A | Qa | - | Dum | - |

**Supplementary Table 8.** Number of molecules in the simulation system. The number of PIP_2_, POPC, Water, Na and Cl and the box size used in the unbiased simulation and the free energy calculation.

| **Protein** | **PIP<sub>2</sub>** | **POPC** | **Water** | **Na** | **Cl** | **Box size** |
| --- | --- | --- | --- | --- | --- | --- |
| hKir1.1 | 1 | 495 | 14196 | 314 | 330 | 134x134x149 Å<sup>3</sup> |
| hKir2.2 | 1 | 538 | 16324 | 378 | 353 | 140x140x155 Å<sup>3</sup> |
| hKir3.2 | 1 | 463 | 12451 | 321 | 285 | 131x131x141 Å<sup>3</sup> |
| hKir6.2 | 1 | 508 | 13989 | 328 | 308 | 135x137x145 Å<sup>3</sup> |

## References

1. Duncan, A. L., Song, W. & Sansom, M. S. P. Lipid-Dependent Regulation of Ion Channels and G Protein–Coupled Receptors: Insights from Structures and Simulations. Annu. Rev. Pharmacol. Toxicol. (2019). doi:10.1146/annurev-pharmtox-010919-023411

2. Gu, R.-X & de Groot, B. L. Lipid-protein interactions modulate the conformational equilibrium of a potassium channel. Nat. Commun. 11, 2162 (2020).

3. Monticelli, L. et al. The MARTINI Coarse-Grained Force Field: Extension to Proteins. J. Chem. Theory Comput. 4, 819 (2008).

4. Marrink, S. J., Risselada, H. J., Yefimov, S., Tieleman, D. P. & de Vries, A. H. The MARTINI Force Field: Coarse Grained Model for Biomolecular Simulations. J. Phys. Chem. B 111, 7812–7824 (2007).

5. Hedger, G. & Sansom, M. S. P. Lipid interaction sites on channels, transporters and receptors: Recent insights from molecular dynamics simulations. Biochim. Biophys. Acta 1858, 2390–2400 (2016).

6. Corey, R. A., Vickery, O. N., Sansom, M. S. P. & Stansfeld, P. J. Insights into Membrane Protein-Lipid Interactions from Free Energy Calculations. J. Chem. Theory Comput. 15, 5727–5736 (2019).

7. Pipatpolkai, T., Usher, S., Stansfeld, P. J. & Ashcroft, F. M. New insights into KATP channel gene mutations and neonatal diabetes mellitus. Nat. Rev. Endocrinol. (2020). doi:10.1038/s41574-020-0351-y

8. Suh, B.-C & Hille, B. PIP2 is a necessary cofactor for ion channel function: how and why? Annu. Rev. Biophys. 37, 175–195 (2008).

9. Rohács, T. et al. Specificity of activation by phosphoinositides determines lipid regulation of Kir channels. Proc. Natl. Acad. Sci. 100, 745 LP–750 (2003).

10. Huang, C.-L, Feng, S. & Hilgemann, D. W. Direct activation of inward rectifier potassium channels by PIP2 and its stabilization by Gβγ. Nature 391, 803–806 (1998).

11. Fan, Z. & Makielski, J. C. Anionic Phospholipids Activate ATP-sensitive Potassium Channels. J. Biol. Chem. 272, 5388–5395 (1997).

12. Hansen, S. B., Tao, X. & MacKinnon, R. Structural basis of PIP2 activation of the classical inward rectifier K+ channel Kir2.2. Nature 477, 495–498 (2011).

13. Whorton, M. R. & MacKinnon, R. Crystal structure of the mammalian GIRK2 K+ channel and gating regulation by G proteins, PIP2, and sodium. Cell 477, 199–208 (2011).

14. Puljung, M. C. Cryo-electron microscopy structures and progress toward a dynamic understanding of KATP channels. J. Gen. Physiol. (2018). doi:10.1085/jgp.201711978

15. Rorsman, P. & Ashcroft, F. M. Pancreatic β-Cell Electrical Activity and Insulin Secretion: Of Mice and Men. Physiol. Rev. (2018). doi:10.1152/physrev.00008.2017

16. De Franco, E. et al. Update of variants identified in the pancreatic β-cell KATP channel genes KCNJ11 and ABCC8 in individuals with congenital hyperinsulinism and diabetes. Hum. Mutat. 41, 884–905 (2020).

17. Haider, S., Tarasov, A. I., Craig, T. J., Sansom, M. S. P. & Ashcroft, F. M. Identification of the PIP2-binding site on Kir6.2 by molecular modelling and functional analysis. EMBO J. 26, 3749–3759 (2007).

18. Reimann, F. et al. Characterisation of new KATP-channel mutations associated with congenital hyperinsulinism in the Finnish population. Diabetologia 46, 241–249 (2003).

19. Wang, L. et al. Accurate and Reliable Prediction of Relative Ligand Binding Potency in Prospective Drug Discovery by Way of a Modern Free-Energy Calculation Protocol and Force Field. J. Am. Chem. Soc. 137, 2695–2703 (2015).

20. Gapsys, V. & de Groot, B. L. pmx Webserver: A User Friendly Interface for Alchemistry. J. Chem. Inf. Model. 57, 109–114 (2017).

21. Gapsys, V., Michielssens, S., Seeliger, D. & de Groot, B. L. pmx: Automated protein structure and topology generation for alchemical perturbations. J. Comput. Chem. 36, 348–354 (2015).

22. Flanagan, S. E. et al. Mutations in ATP-sensitive K+ channel genes cause transient neonatal diabetes and permanent diabetes in childhood or adulthood. Diabetes 56, 1930–1937 (2007).

23. Huopio, H. et al. Acute Insulin Response Tests for the Differential Diagnosis of Congenital Hyperinsulinism. J. Clin. Endocrinol. Metab. 87, 4502–4507 (2002).

24. Stansfeld, P. J. et al. MemProtMD: Automated Insertion of Membrane Protein Structures into Explicit Lipid Membranes. Structure 23, 1350–1361 (2015).

25. Fajer, M., Swift, R.V & McCammon, J. A. Using multistate free energy techniques to improve the efficiency of replica exchange accelerated molecular dynamics. J. Comput. Chem. 30, 1719–1725 (2009).

26. Bonomi, M. et al. PLUMED: A Portable Plugin for Free-Energy Calculations with Molecular Dynamics. Comput. Phys. Commun. 180, 1961 (2009).

27. Tribello, G. A., Bonomi, M., Branduardi, D., Camilloni, C. & Bussi, G. PLUMED 2: New feathers for an old bird. Comput. Phys. Commun. 185, 604–613 (2014).

28. Duncan, A. L., Corey, R. A. & Sansom, M. S. P. Defining how multiple lipid species interact with inward rectifier potassium (Kir2) channels. Proc. Natl. Acad. Sci. 117, 7803 LP–7813 (2020).

29. Trapp, S., Proks, P., Tucker, S. J. & Ashcroft, F. M. Molecular Analysis of ATP-sensitive K Channel Gating and Implications for Channel Inhibition by ATP. J. Gen. Physiol. 112, 333 LP–349 (1998).

30. Schwappach, B., Zerangue, N., Jan, Y. N. & Jan, L. Y. Molecular Basis for K_ATP_ Assembly: Transmembrane Interactions Mediate Association of a K^+^ Channel with an ABC Transporter. Neuron 26, 155–167 (2000).

31. Schacht, J. Purification of polyphosphoinositides by chromatography on immobilized neomycin. J. Lipid Res. 19, 1063–1067 (1978).

32. Peter, P. et al. Running out of time: the decline of channel activity and nucleotide activation in adenosine triphosphate-sensitive K-channels. Philos. Trans. R. Soc. B Biol. Sci. 371, 20150426 (2016).

33. Tucker, S. J., Gribble, F. M., Zhao, C., Trapp, S. & Ashcroft, F. M. Truncation of Kir6.2 produces ATP-sensitive K+ channels in the absence of the sulphonylurea receptor. Nature 387, 179–183 (1997).

34. Baukrowitz, T. et al. PIP2 and PIP as determinants for ATP inhibition of KATP channels. Science (80-.). 282, 1141–1144 (1998).

35. Domański, J., Hedger, G., Best, R. B., Stansfeld, P. J. & Sansom, M. S. P. Convergence and Sampling in Determining Free Energy Landscapes for Membrane Protein Association. J. Phys. Chem. B 121, 3364–3375 (2017).

36. Hedger, G. et al. Lipid-Loving ANTs: Molecular Simulations of Cardiolipin Interactions and the Organization of the Adenine Nucleotide Translocase in Model Mitochondrial Membranes. Biochemistry 55, 6238 (2016).

37. Naughton, F. B., Kalli, A. C. & Sansom, M. S. P. Modes of Interaction of Pleckstrin Homology Domains with Membranes: Toward a Computational Biochemistry of Membrane Recognition. J. Mol. Biol. 430, 372–388 (2018).

38. Webb, B. & Sali, A. Comparative Protein Structure Modeling Using MODELLER. Curr. Protoc. Bioinformatics 54, 5.6.1–5.6.37 (2016).

39. Martin, G. M., Kandasamy, B., DiMaio, F., Yoshioka, C. & Shyng, S.-L Anti-diabetic drug binding site in a mammalian KATP channel revealed by Cryo-EM. Elife 6, e31054 (2017).

40. Waterhouse, A. et al. SWISS-MODEL: homology modelling of protein structures and complexes. Nucleic Acids Res. 46, W296–W303 (2018).

41. Newport, T. D., Sansom, M. S. P. & Stansfeld, P. J. The MemProtMD database: a resource for membrane-embedded protein structures and their lipid interactions. Nucleic Acids Res. 47, D390–D397 (2018).

42. Nugent, T. & Jones, D. T. Membrane protein orientation and refinement using a knowledge-based statistical potential. BMC Bioinformatics 14, 276 (2013).

43. Bussi, G., Donadio, D. & Parrinello, M. Canonical sampling through velocity rescaling. J. Chem. Phys. 126, 14101 (2007).

44. Parrinello, M. & Rahman, A. Polymorphic transitions in single crystals: A new molecular dynamics method. J. Appl. Phys. 52, 7182–7190 (1981).

45. Abraham, M. J. et al. Gromacs: High performance molecular simulations through multi-level parallelism from laptops to supercomputers. SoftwareX 1-2, 19–25 (2015).

46. Schrodinger LLC. The PyMOL Molecular Graphics System, Version 1.8. (2015).

47. Klimovich, P. V, Shirts, M. R. & Mobley, D. L. Guidelines for the analysis of free energy calculations. J. Comput. Aided. Mol. Des. 29, 397–411 (2015).

48. Hub, J. S., De Groot, B. L. & Van Der Spoel, D. g_wham-A Free Weighted Histogram Analysis Implementation Including Robust Error and Autocorrelation Estimates. J. Chem. Theory Comput. 6, 3713 (2010).

49. Proks, P., Girard, C., Baevre, H., Njolstad, P. R. & Ashcroft, F. M. Functional effects of mutations at F35 in the NH2-terminus of Kir6.2 (KCNJ11), causing neonatal diabetes, and response to sulfonylurea therapy. Diabetes 55, 1731–1737 (2006).

